# H2A.Z levels control the timing of major events at the maternal-zygotic transition

**DOI:** 10.64898/2026.04.27.721027

**Authors:** Pakinee Phromsiri, Xiaolu Wei, Claire E. Makowski, Noah Reger, Duy K. Nguyen, Humaira Marzia Alam, Yuki Shindo, Amanda Amodeo, Patrick J. Murphy, Fanju W. Meng, Michael A. Welte

**Affiliations:** Department of Biology, University of Rochester, Rochester, NY, USA; Department of Biological Sciences, University of North Texas, Denton, TX, USA; Department of Biomedical Genetics, University of Rochester Medical Center, Rochester, NY, USA; Department of Pathology, University of Rochester Medical Center, Rochester, NY, USA; Department of Molecular Biology and Genetics, Cornell University, Ithaca, NY, USA; Department of Biological Sciences, Dartmouth College, Hanover, NH, USA

## Abstract

Across animals, early embryonic events are temporally tightly coordinated but the underlying mechanisms remain incompletely understood. In Drosophila, both global and nuclear levels of the histone variant H2Av rise progressively during the maternal-zygotic transition (MZT), but whether this increase is functionally important is unknown. We find that increased *H2Av* dosage expedites specific MZT events: the transition from nuclear cycle (NC) 13 to 14 occurs prematurely as does the turnover of thousands of maternal transcripts; in addition, a subset of genes is precociously expressed in the zygote. Reduced *H2Av* dosage has reciprocal effects. Comparable transcriptional shifts are observed in zebrafish embryos overexpressing the H2Av ortholog H2A.Z, suggesting *H2Av/H2A.Z* dosage as a conserved timer of early development. We also examined mutants with impaired H2Av sequestration on lipid droplets which exhibit increased nuclear H2Av levels but reduced cytoplasmic levels. Unexpectedly, nuclear H2Av abundance influences the timing of NC 13 but is not the main driver of transcriptome remodeling. In summary, we find that H2Av/H2A.Z levels are critical timers of early embryogenesis and that H2Av can act in part via a non-nuclear mechanism.

**Author summary:** In animals, the earliest stages of embryogenesis are initially driven by proteins and RNAs that the mother provides via the egg; later development is controlled by the embryo’s own genes. The switch from maternal to zygotic control is called the maternal-zygotic transition (MZT). This conserved process involves degradation of maternal mRNAs, activation of zygotic genes, cell cycle lengthening, and morphological remodeling. All these events require precise control, but how they are coordinated remains incompletely understood. Here we show that the levels of a specific histone, H2A.Z, provide a timer for the MZT. In Drosophila, H2A.Z levels – both in the embryo overall and in nuclei – increase during the MZT. When we ectopically increased H2A.Z levels, specific MZT events occurred prematurely. Reducing H2A.Z levels had the opposite effect. Using mutant flies in which H2A.Z levels are ectopically increased in the nucleus but reduced in the embryo as a whole, we uncovered both nuclear and non-nuclear roles for H2A.Z in the establishment of developmental timing. Moreover, we observed similar outcomes in genetically manipulated zebrafish embryos. Our results are consistent with an ancient mechanism in which H2A.Z abundance functions as a timer of early embryogenesis.

## Introduction

Metazoans typically develop from a single-celled zygote loaded with maternally supplied factors that drive the earliest stages of development in the absence of transcription; this maternal control is eventually replaced by zygotic transcription, which becomes the dominant regulator of development. The handoff from maternal to zygotic control is part of a process called the maternal-to-zygotic transition (MZT) and is vital for the activation of developmental programs, including pluripotency, morphogenesis, and differentiation. (1–3). Despite extensive study, however, the regulatory mechanisms underlying this switch remain incompletely understood.

*Drosophila melanogaster* provides a powerful model to investigate this switch. For the first ∼2.5 hours after fertilization, Drosophila embryos undergo a series of 13 rapid, near synchronous mitotic divisions without cytokinesis, resulting in an embryo with around 6000 diploid nuclei (4); the period from one mitosis to the next is called a nuclear cycle (NC). These early divisions do not require zygotic transcription (5–7). The MZT begins around NC 8, when minor zygotic genome activation (ZGA) occurs, which results in limited transcription of early zygotic genes. This minor wave is accompanied by a progressive lengthening of cell cycles. A critical developmental switch occurs in NC 13 and 14: the major wave of ZGA is initiated, leading to robust transcription of a broad array of zygotic genes (1). During the MZT, the embryo also undergoes dramatic morphological changes: the peripheral cytoplasm becomes transparent due to inward transport of various yolk organelles (cytoplasmic clearing), then thousands of cells form in this region (cellularization), followed by gastrulation movements. The remodeling of the transcriptome, cell cycle lengthening, and morphological changes are highly coordinated and occur in a consistent temporal pattern. However, the molecular mechanisms that regulate the timing of these MZT events remain incompletely understood.

Previous studies have identified the nuclear-to-cytoplasmic (N:C) ratio as a critical regulator for MZT timing, based on the observation that increasing nuclear DNA content relative to a fixed cytoplasmic volume can trigger ZGA once a threshold is reached (5, 8–10). More recent work has revealed that N:C-dependent mechanisms control the timing of a specific subset of MZT events and that the N:C ratio is largely mediated through the availability of canonical histones (11–16).

In early embryos, the levels of both canonical and variant histones are temporally tightly controlled. The pre-MZT embryo requires large amounts of histones because the amount of its DNA doubles every 8-20 minutes. This need is partially met by histone stores that accumulate in the oocyte cytoplasm during oogenesis; these stores are further supplemented by the translation of maternally deposited mRNAs so that histone protein levels go up globally. This pattern has been observed both for canonical (17, 18) and variant histones (17). However, while the nuclear levels of canonical histones H2A, H2B, and H3 either stay constant or decrease during NC 10-NC 13 (17, 19), they go up for the variant histones H2Av and H3.3. (17, 19, 20). Thus, the nuclear levels of histones are temporally controlled during early embryogenesis and might – in principle – mediate the timing of events of the MZT. However, the functional importance of this regulation is unknown.

Drosophila H2Av combines the functions of H2A.Z and H2A.X in other organisms (21). Similar to H2A.X, H2Av is phosphorylated in response to DNA double-strand breaks, promoting DNA damage signaling and repair processes. Akin to H2A.Z, it is enriched at promoters and enhancers, where it modulates transcriptional activity. Both functions are already evident in embryos during MZT: H2Av is phosphorylated in response to irradiation (22), and dramatic reduction of chromatin-bound H2Av (in response to knockdown of the H2Av chaperone Domino) results in reduced transcription of hundreds of genes (23). Together, these observations point to a central role for H2Av in coordinating genome integrity with transcriptional activation during early embryogenesis.

Intriguingly, prior to the MZT, nuclear accumulation of H2Av is not determined solely by protein abundance but is also regulated by cytoplasmic sequestration. H2Av associates dynamically with the anchoring protein Jabba on lipid droplets, abundant fat storage organelles in the cytosol. This interaction slows H2Av import into nuclei, leading to lower nuclear H2Av levels than without sequestration (17, 20). The existence of such a mechanism may suggest that achieving particular H2Av levels in the nucleus is functionally important, but that notion has not yet been critically tested.

Here we investigate the role of H2Av in regulating MZT timing. We compare embryos with varying *H2Av* gene dosages and monitor their developmental progression by live imaging and their transcriptomes by RNA sequencing. The timing of NC 13 is highly sensitive to *H2Av* dosage; mitosis 13 is expedited in embryos with increased *H2Av* dosage and delayed with decreased H2Av levels. Embryos with increased *H2Av* dosage start out with a similar set of transcripts as the wild type but by NC 13 about one-sixth of transcripts are significantly altered, with most of them displaying an expedited developmental trajectory. Embryos with reduced *H2Av* dosage exhibit the opposite pattern. Using mutants with elevated nuclear H2Av levels but reduced cytoplasmic levels, we find that nuclear H2Av abundance influences timing of NC 13 but is not the main driver of transcriptome remodeling. Finally, to determine evolutionary conservation, we examine the developmental trajectory of the transcriptome in zebrafish embryos overexpressing the H2Av ortholog H2A.Z. Many transcripts mimicked the pattern observed in Drosophila, with precocious up– or downregulation, indicating conserved function. We conclude that H2Av/H2A.Z levels are critical timers of early embryogenesis in both Drosophila and zebrafish.

## Results

### H2Av levels time nuclear cycle length

During NC 11 to NC 14, the nuclear levels of H2Av increase (17, 20) while those of canonical H2A decrease (Figure S1A, S1B). Thus, H2Av accumulation within the nucleus tracks with developmental time, but it remains unknown whether differential H2Av levels causatively affect the timing of development. We therefore generated genotypes expressing either one, two, or four copies of *H2Av*, confirmed protein expression levels, and examined the developmental progression of their embryos.

To monitor H2Av dynamics *in vivo*, we tagged H2Av at its C-terminus with the fluorescent protein Dendra2. Flies homozygous for this allele (*Endo-H2Av-Dendra2*) served as our wild-type comparison and will hereafter be referred to as *WT H2Av*. To achieve increased H2Av levels, we introduced a previously characterized *H2Av-Dendra2* (*gH2Av*) transgene (20) into this genetic background; this transgene carries the *H2Av* locus (including the promoter region and 3’UTR) with a C-terminal Dendra2 tag. Flies homozygous for both *Endo-H2Av-Dendra2* and the transgene are referred to as *Double H2Av*. To create a condition with reduced *H2Av* expression, flies heterozygous for *Endo-H2Av-Dendra2* and the null allele *H2Av^KO^* (17, 24) were used; this genotype will hereafter be referred to as *Half H2Av*. Embryos analyzed were derived from mothers of this genotype crossed to wild-type fathers; while this cross generates several zygotic genotypes, we expect that H2Av levels are largely determined by the maternal genotype, as *H2Av* mRNA and protein are maternally provided in large amounts (17, 20).

As expected, global H2Av protein levels in *Half H2Av* embryos were lower compared to *WT H2Av*, while levels were elevated in *Double H2Av* embryos (Figure 1A, 1B). The same pattern was seen for nuclear H2Av levels (Figure 1C, 1D). For *Half H2Av* and *WT H2Av* embryos, nuclear H2Av levels increased developmentally (Figure S1C,D), as previously observed for wild-type embryos without the Dendra2 tag (17); for Double H2Av, nuclear H2Av levels in NC 11 and NC 12 were already at NC 13 levels (Figure S1E). Thus, if nuclear H2Av levels are important to trigger specific downstream developmental events, those should occur earlier in *Double H2Av* embryos and later in *Half H2Av*, compared to *WT H2Av* embryos.

**Figure 1.**
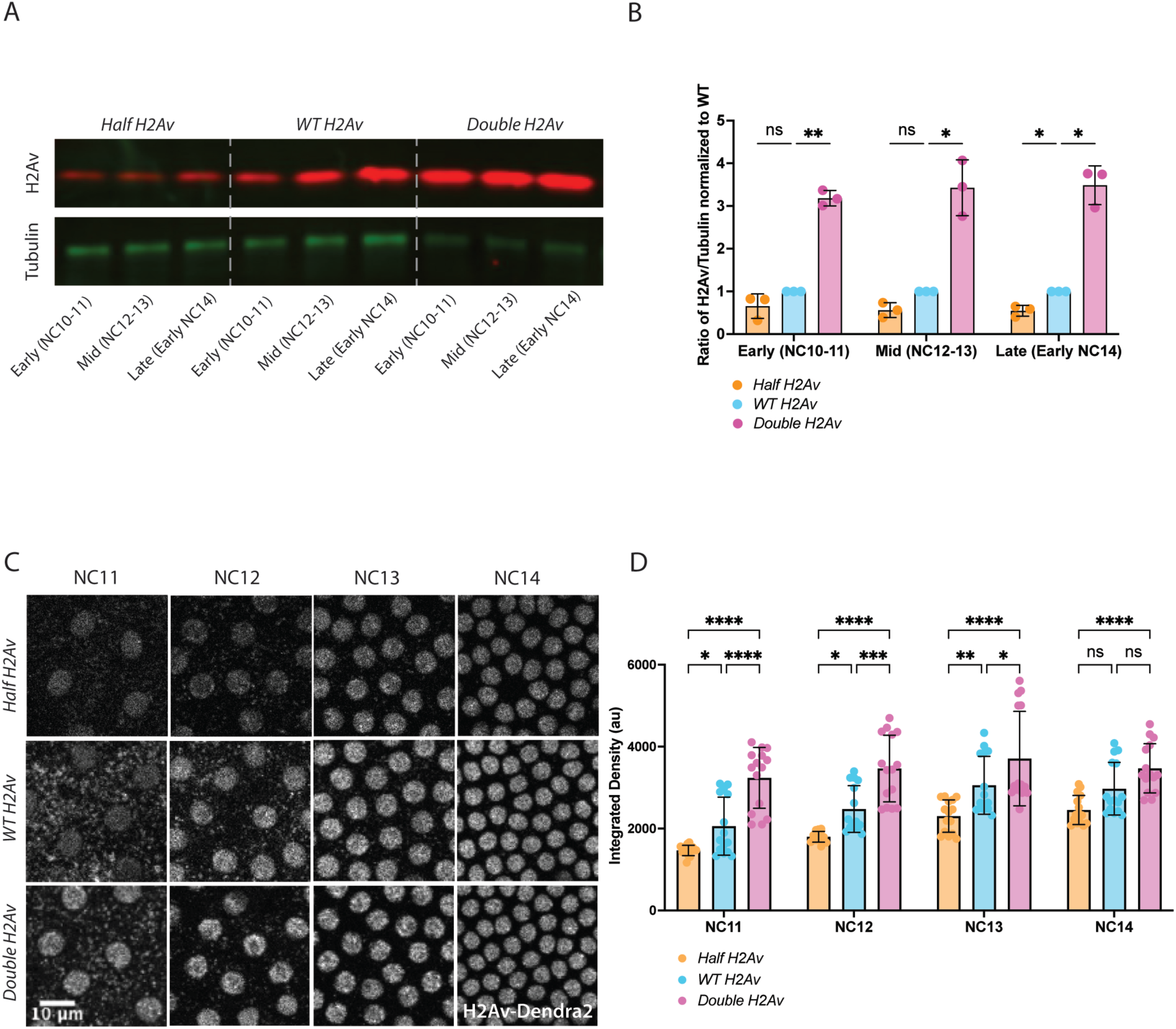
Nuclear H2Av increases developmentally, and protein levels can be modulated via altering H2Av gene dosage. (A) Total H2Av protein levels correlate with gene dosage. Protein extracts from 10 visually staged embryos per condition (*Half H2Av*, *WT H2Av*, and *Double H2Av*) at Early (NC 10-NC 11), Mid (NC 12-NC 13), and Late (pre-cellularization NC 14) blastoderm were analyzed by SDS-PAGE and Western blotting. Membranes were probed with antibodies against H2Av (red) as well as against alpha-tubulin (green), as a loading control. (B) Quantitation of Western blots like in (A) for *Half H2Av*, *WT H2Av*, and *Double H2Av* embryos normalized to *WT H2Av* signal at each corresponding stage. Error bars represent standard deviation (n=3). Statistical analysis was performed using two-way ANOVA followed by Tukey’s multiple comparisons test. (C) Representative z-projection images of embryos during interphase of NC 11, 12, 13, and 14, expressing one, two, or four copies of *H2Av-Dendra2* (referred to as *Half H2Av*, *WT H2Av*, and *Double H2Av*, respectively). Images show increased nuclear H2Av-Dendra2 accumulation with increasing gene dosage. Scale bar: 10 µm. (D) Quantitation of total nuclear H2Av-Dendra2 fluorescence intensity per nucleus during interphase of NC 11-NC 14 in *Half H2Av*, *WT H2Av*, and *Double H2Av* embryos. Five nuclei were measured per embryo for three embryos per genotype. Error bars represent mean with SEM. Statistical analysis was performed using two-way ANOVA followed by Tukey’s multiple comparisons test. * for P ≤ 0.05, ** for P ≤ 0.01, *** for P ≤ 0.001, and **** for P ≤ 0.0001, “ns” = not significant (P ≥ 0.05).

Embryos of all three genotypes developed grossly normally, progressing through syncytial stages, cellularization, gastrulation, and germband extension; they all hatched at high rates. However, when we imaged the embryos live, we observed frequent formation of chromatin bridges during syncytial mitoses in *Half* and *Double H2Av* embryos (Figure S2A), indicative of under-replicated or mis-segregated DNA (25, 26). Consistent with DNA damage, we observed increased nuclear falling, an embryo-specific response where damaged nuclei at the embryo surface migrate towards the central region (26) (Figure S2B). Thus, while altered H2Av levels do not impair overall developmental progression, they do induce Chk2-dependent DNA damage–associated phenotypes (27, 28).

To determine if the timing of developmental progression was affected, we performed live confocal imaging of embryos progressing from NC 10 to NC 14 at 25°C. These cell cycles are tightly controlled temporally and consistent from embryo to embryo when environmental conditions are kept constant. As H2Av-Dendra2 is present on chromatin and highlights the mitotic chromosomes, our videos allowed us to monitor when mitosis occurred (Figure 2A, 2B). Counting nuclear cycle length from one anaphase to the next, we observed the known gradual lengthening of the cell cycle from NC 11 to NC 13 (Figure 2A, B; Figure S3A, S3B) in *WT H2Av* embryos. Under our conditions, this represents an increase from 10.9 to 19.0 minutes. The length of NC 13 was altered in *Half H2Av* and *Double H2Av* embryos. Increased *H2Av* dosage expedited entry into mitosis 13 by ∼8%, and reduced *H2Av* dosage delayed NC 13 completion by ∼5%. Given how tightly controlled these cell cycles usually are, these timing changes were substantial, well outside wild-type variation ranges, and found to be statistically significant. We conclude that H2Av levels have a specific effect on regulating the timing of mitosis 13, with lower levels delaying the entry into mitosis 13 and higher levels expediting it. Interestingly, all three genotypes had very similar durations for NC 11 and NC 12 (Figure S3A, S3B). Thus, the delayed or expedited entry into mitosis 13 does not appear to reflect an overall slow-down or speed up of early development but a temporally specific effect.

**Figure 2.**
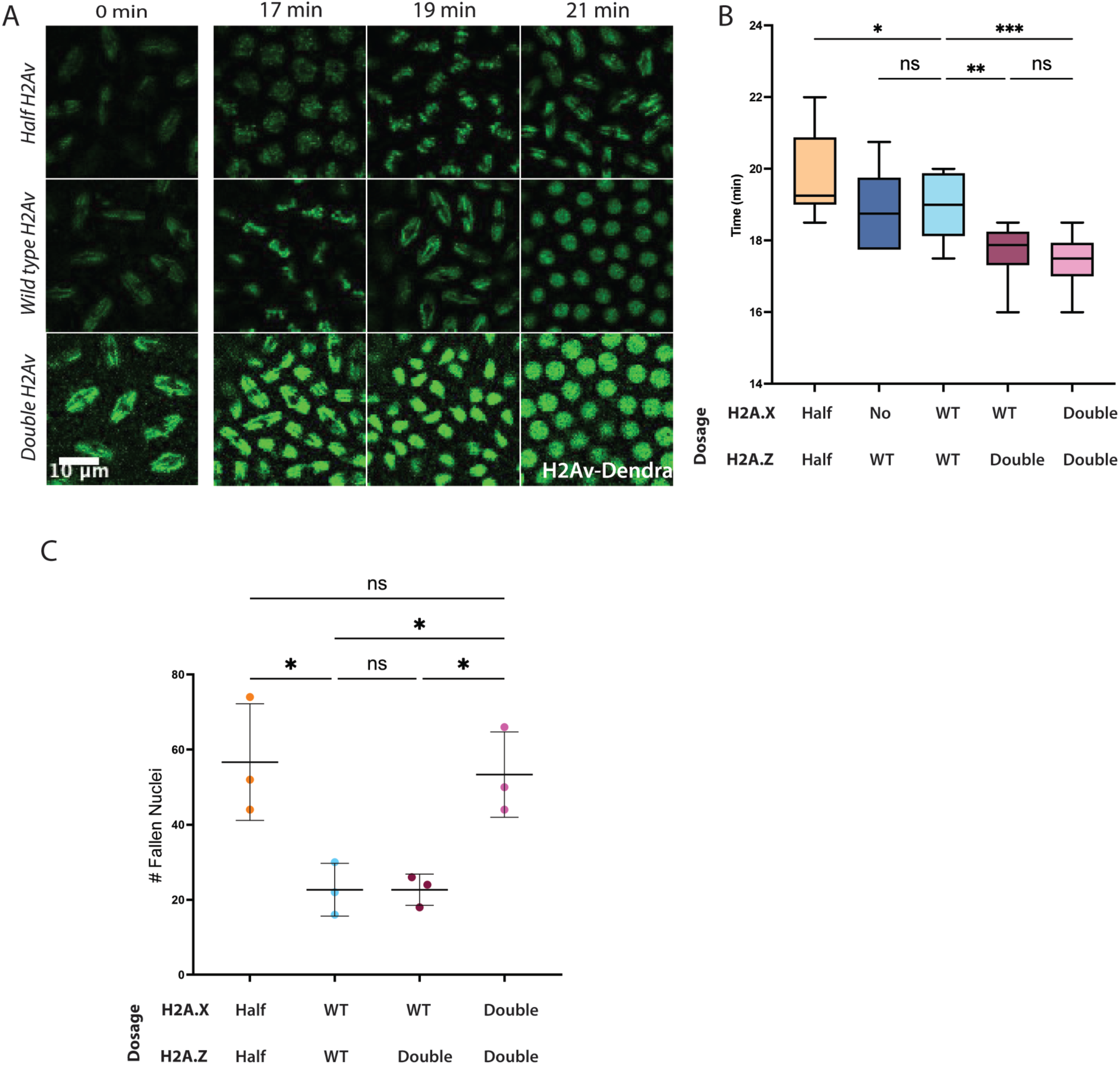
*H2Av* dosage controls NC 13 timing and H2A.X function is dispensable for NC 13 timing but promotes nuclear falling. *Half H2Av*, *WT H2Av,* and *Double H2Av* embryos were imaged at 25°C during NC 10-NC 14. The duration of nuclear cycles was quantified from time-lapse movies by measuring the interval between anaphase onset of one mitosis and the next. (A) Still frames from representative live-imaging movies showing progression through NC 13 in *Half H2Av*, *WT H2Av*, and *Double H2Av* embryos. NC 13 is ∼1 minute longer in *Half H2Av* embryos and ∼1.5 minutes shorter in *Double H2Av* embryos compared to *WT H2Av*. Each violin plot displays the full distribution with a line at the median. N=3 embryos per genotype. (B) Box plots showing NC 13 duration in genotypes with varying *H2A.Z* and *H2A.X* dosage. Line at median. p values were calculated using ordinary one-way ANOVA followed by uncorrected Fisher’s LSD. ****=p<.0001. N=11-12 embryos. (C) Box plot showing quantitation of the number of fallen nuclei in NC 14 embryos. The degree of nuclear falling is increased in embryos *Double H2A.Z*, *Double H2A.X* embryos compared to *Double H2A.Z*, *WT H2A.X* embryos. Reduction of H2A.X in *Double H2A.Z, WT H2A.X* rescued NF to near *WT* conditions. Reduction of H2Av (*Half H2A.Z, Half H2A.X*) results in increased NF. P values were calculated using one-way ANOVA followed by Tukey’s test. * for P ≤ 0.05, ** for P ≤ 0.01, *** for P ≤ 0.001, and **** for P ≤ 0.0001, “ns” = not significant (P ≥ 0.05) N = 5-6.

### Phosphorylation of Ser137, and thus H2A.X function, is dispensable for nuclear cycle timing

H2Av has a dual nature, combining the functions of H2A.Z and H2A.X in other organisms (21). Like H2A.Z, it contributes to transcriptional regulation and chromatin structure, and like H2A.X, it mediates DNA damage responses, via phosphorylation at serine137. Either of these functions could contribute to the altered NC 13 duration and nuclear falling. NC 13 duration is known to be regulated by the DNA damage checkpoint, with embryos lacking ATR or Chk1 exhibiting expedited mitosis (29, 30), and we found that nuclei undergoing fallout were reproducibly positive for the phosphorylated form of H2Av (Figure S2C, S2D). These observations are consistent with a model that H2A.X function is responsible for both nuclear falling and the timing of mitosis 13.

To investigate this idea, we generated a genomic *H2Av* transgene tagged with Dendra2 in which the codon for serine137 was mutated to encode alanine (Figure S3C). This change should render H2Av non-phosphorylatable and thus unable to mediate the DNA repair-associated activity of H2A.X. We will refer to this transgene as *H2Av^S137A^-Dendra2*. Embryos from mothers expressing *H2Av^S137A^– Dendra2* either in the absence or presence of wild-type H2Av at the endogenous locus were viable and developed grossly normally; thus, H2A.X function is not required for embryogenesis per se. Using this transgene, it was now possible to increase the dosage of *H2A.Z* without increasing the dosage of *H2A.X*, thereby allowing us to distinguish which embryonic phenotypes were affected by the H2A.X-like or H2A.Z-like functions of H2Av.

We first examined embryos expressing two copies of *H2Av^S137A^-Dendra* in the presence of wild-type H2Av at the endogenous locus. Based on our hypothesis, these embryos should have normal H2A.X function but doubled H2A.Z activity (*WT H2A.X, Double H2A.Z*). Unlike in *Double H2Av* (*Double H2A.X, Double H2A.Z*) embryos, levels of nuclear falling were not increased and were similar to those in *WT H2Av* (Figure 2C), indicating that increased dosage of *H2A.Z* did not trigger the DNA damage response. This result also implies that the increased nuclear falling in *Double H2Av* embryos must be due to excess H2A.X. In contrast, these *WT H2A.X, Double H2A.Z* embryos displayed a shorter NC 13 length compared to *WT H2Av*, indicating that increasing *H2A.Z* dosage is sufficient to expedite mitosis 13, similar to embryos with four copies of *WT H2Av* (*Double H2A.X, Double H2A.Z)* (Figure 2B). Thus, the effect of H2Av on cell-cycle timing can be decoupled from its role in DNA damage signaling.

We next analyzed embryos expressing two copies of *H2Av^S137A^-Dendra2* in the absence of endogenous H2Av. These embryos should have normal H2A.Z function but lack H2A.X function (*No H2A.X, WT H2A.Z*). NC 13 timing was comparable to embryos that express two copies of wild-typ*e H2Av* (*WT H2A.X, WT H2A.Z*); therefore, loss of the H2A.X-like function did not result in the delayed mitosis 13 seen in *Half H2Av* embryos (Figure 2B).

Taken together, these results indicate that phosphorylation of H2Av at serine137, and by extension the H2A.X function in DNA damage sensing, is important for nuclear falling but dispensable for the regulation of NC 13 timing. In contrast, the *H2A.Z*-like dosage does not affect nuclear falling but alters the regulation of NC 13 timing. Our data thus indicate that the effects on NC 13 timing and nuclear falling can be uncoupled, arguing that nuclear falling does not cause the observed change in NC 13 or vice versa.

### Elevated H2A.Z levels result in changes to one-sixth of the transcriptome by the ZGA

We next investigated whether H2Av levels specifically affect the timing of mitosis 13 or also that of other developmental events. H2A.Z has established roles in modulating transcription (31–34), and in Drosophila and zebrafish embryos, H2Av (H2A.Z in fish) is necessary for proper transcription of many genes at ZGA (23, 35–37). Nuclear accumulation of H2Av increases progressively (Figure 1B, S1D) from NC 11 to NC 13. If the transcription of a particular H2Av target gene is triggered when a certain threshold of H2Av is reached, its expression level should be dependent on *H2Av* dosage.

We therefore investigated the transcriptome of *Double H2Av* and *WT H2Av* embryos at three timepoints: shortly after egg laying (referred to as NC 1 going forward), in NC 13, and in NC 14. For NC 13 and NC 14, we visually staged embryos expressing H2Av-Dendra2 via live imaging; embryos were allowed to progress through mitosis 11 or 12, recovered 8 or 15 min later, respectively, and then flash frozen individually. cDNAs prepared from isolated mRNA samples were then sequenced for each sample (Figure S4A), and differential expression analysis was conducted using DESeq2. The NC 1 timepoint represents solely transcripts that are maternally provided. By the NC 13 and NC 14 timepoints, a large fraction of the maternal transcripts is known to be supplemented by zygotic transcription (38, 39); thus, the NC 13 and NC 14 samples should represent such maternal-zygotic transcripts as well as purely maternal and purely zygotic ones.

To establish a baseline for comparison between genotypes, we first characterized how in *WT H2Av* embryos the transcriptome changes as the embryos develop. RNA transcripts detectible during MZT could result from maternal deposition or zygotic gene transcription. For the sake of clarity, we therefore refer to a collection of identical RNA transcripts as a “transcriptional unit”, rather than an expressed gene. Our samples showed an increase in total number of detectable transcriptional units over developmental time, with 7540 transcriptional units for NC 1, 9805 transcriptional units for NC 13, and 10594 transcriptional units for NC 14. This pattern is consistent with the maternally provided transcripts being supplemented by zygotic transcription by NC 13 and NC 14. Those transcriptional units absent in NC 1, but detected at NC 13 and NC 14, must be due to zygotic transcription, and we will refer to them as “zygotic transcripts” going forward. They accounted for 2411 transcriptional units at NC 13 (products of the minor wave of ZGA) and 3169 at NC 14 (products of both the minor and major wave). Of the 2411 zygotic transcripts detectable at both stages, their expression either did not change significantly from NC 13 to NC 14 (gray dots) or was robustly upregulated (blue dots) (Figure S4C’). Thus, both the number and levels of zygotic transcripts increased from NC 13 and NC 14, consistent with our analysis detecting the major wave of ZGA.

The temporal trajectory of maternally provided transcriptional units is expected to be more complicated; as embryogenesis proceeds, they may be degraded but also replenished by zygotic transcription (38–40). As the literature has not yet arrived at a consensus which of the transcripts deposited by the mother into the egg are supplemented by zygotic transcription (“maternal-zygotic” transcripts) and which ones are not (“exclusively maternal transcripts”), here we use the term maternal transcriptional units for those that are detectable in NC 1 embryos. From NC 1 to NC 13, most of these maternal transcriptional units did not show a statistically significant change in levels during this period (gray dots); 874 decreased in levels and 791 increased (Figure S4B). From NC 13 to NC 14, the overall trend is similar, with many transcriptional units unchanged, but 1931 net decreased and 2265 increased (Figure S4C). For the increasing transcriptional units, new production must outweigh the loss through RNA degradation; for the decreasing ones, degradation predominates.

Using this general framework, we next compared the transcriptomes of *Double H2Av* and *WT H2Av* embryos. For the NC 1 samples, the transcriptomes of the two genotypes were largely indistinguishable, with only ∼4.5% of transcriptional units exhibiting differential expression (Figure S4D); 137 of them were significantly lower and 164 significantly higher (Figure S4G). However, by NC 13, expression of more than 17% of the transcriptome (1929 transcriptional units, total) was significantly altered (Figure S4E-E’), and at NC 14, about 16% was altered (1782 transcriptional units) (Figure S4F-F’). Thus, the mothers of the two genotypes largely endow their embryos with similar levels of transcripts for most genes but as development proceeds the transcriptomes diverged.

We first focused on those transcriptional units that were expressed at NC 13 and/or NC 14 but undetectable in NC 1 embryos. Surprisingly, these purely zygotic transcripts were mostly unaffected in *Double H2Av* compared to *WT H2Av* embryos. At NC 13, only 79 of 4395 such transcripts (1.8%) were differentially expressed (Figure S4E’); 35 were downregulated and 44 were upregulated in *Double H2Av* embryos compared to WT (Figure S4H’). At NC 14, only 84 (1.9%) displayed significant changes (Figure S4F’), with 54 lower and 30 higher in *Double H2Av* embryos (Figure S4I’). Thus, strictly zygotic transcripts at ZGA were only modestly impacted by increased H2Av levels, with a slight bias for upregulation at NC 13 and for downregulation at NC 14. A possible explanation for this modest effect on zygotic transcription would be that if the extra H2Av in the nuclei of *Double H2Av* embryos is not incorporated into chromatin but simply present in the nucleoplasm. To address this issue, we analyzed our videos of developing embryos to quantify the H2Av-Dendra2 signal on mitotic chromosomes during metaphase. This chromosome-associated H2Av was roughly double in *Double H2Av* compared to *WT H2Av* embryos (Figure S9), strongly indicating that *Double H2Av* embryos have indeed significantly increased levels of H2Av in chromatin.

Our observations suggest that most of the differences between the two genotypes were due to genes whose transcripts are maternally provided and reflect how rapidly these transcriptional units are degraded in the embryos and/or how much they are replenished by zygotic transcription. To minimize background signal from very low-abundance transcripts, we restricted our analysis to genes with transcripts per million (TPM)> 1 (6661 out of 7540). Only 301 (∼4.5%) showed differential maternal loading in NC 1 embryos (Figure S4D). In contrast, 1850 (∼28 %) were differentially expressed at NC 13 (Figure S4E), with 1141 significantly downregulated and 709 significantly upregulated (Figure S4H), and 1698 (∼25 %) at NC 14 (Figure S4F), with 1173 significantly downregulated and 525 significantly upregulated (Figure S4I). Thus, the majority of differences between the two genotypes were observed for transcriptional units already detected in NC 1 embryos, *i.e.*, purely maternal and maternal/zygotic transcripts. Of those maternally provided transcriptional units, about a quarter were differentially expressed by the ZGA. Together, these data show that increased H2Av dosage alters a substantial fraction of the embryonic transcriptome by the time of ZGA, with effects primarily concentrated on transcriptional units with a maternal contribution rather than newly activated purely zygotic genes.

### Elevated H2Av levels expedite the onset of maternal-to-zygotic transcriptome remodeling while lowered H2Av levels delay it

Clearly, increased H2Av levels altered the levels of many transcriptional units by NC 13/NC 14, predominately, those which were maternally provided. But whether the observed expression changes were indicative of an altered developmental time course or representative of misregulation unrelated to developmental progression remained unclear. One possibility is that these changes simply represent aberrant transcription of maternal genes that are usually zygotically silent. To determine if there is a consistent temporal pattern to the observed changes, we next compared the transcript dynamics in *WT H2Av* and *Double H2Av* embryos at the ZGA.

To track developmental progression, we computed the Log2Fold change (Log2FC) between NC 13 and NC 14 in *WT H2Av* embryos. Against this developmental axis, we plotted the Log2FC in transcript levels of *Double H2Av* vs *WT H2Av* at the three timepoints to capture the effect of the genotype: NC 1 (Figure 3B), NC 13 (Figure 3C), and NC 14 (Figure 3D). This allowed us to classify the transcriptional units into four categories, as outlined in Figure 3A. Quadrant I (QI) transcriptional units were upregulated at the ZGA (from NC 13 to NC 14) and increased in *Double H2Av* embryos relative to *WT H2Av* (positive developmental Log2FC, positive genotype Log2FC). Quadrant II (QII) transcriptional units were normally downregulated at ZGA and increased in *Double H2Av* embryos (negative developmental Log2FC, positive genotype Log2FC). Quadrant III (QIII) transcriptional units were normally downregulated at ZGA and lower in *Double H2Av* vs *WT H2Av* embryos (negative developmental Log2FC, negative genotype Log2FC). Quadrant IV (QIV) transcriptional units were upregulated developmentally but lower in *Double H2Av* embryos compared to *WT H2Av* (positive developmental Log2FC, negative genotype Log2FC).

**Figure 3.**
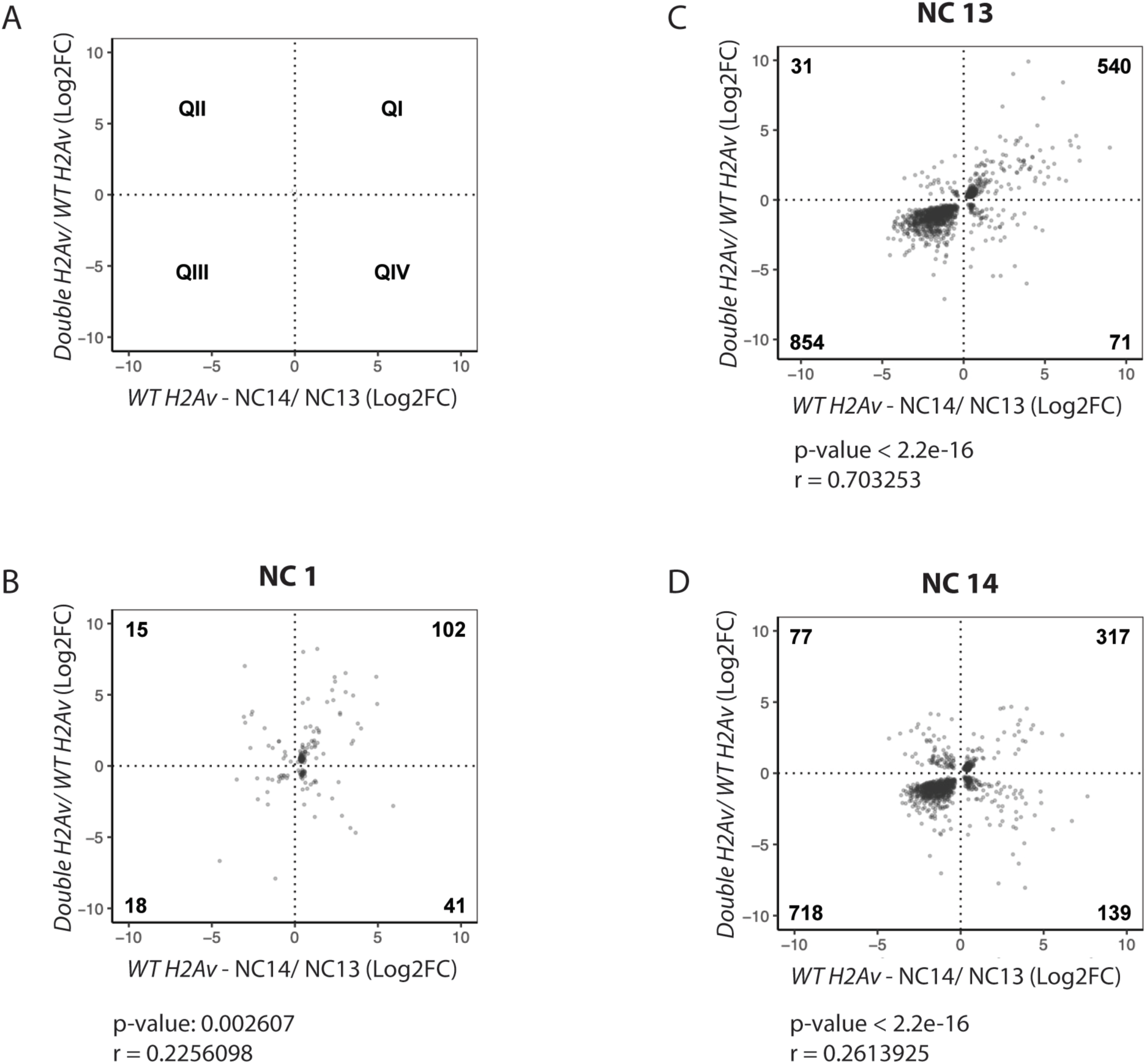
Elevated H2Av levels expedites transcriptional landscape at ZGA. Comparison of DEGs in *Double H2Av* compared to *WT H2Av* during the ZGA (NC 13-NC 14) reveals a subset of transcriptional units that are downregulated prematurely and upregulated precociously. (A) Transcriptional units were plotted based on their Log2FC (p<0.05) during ZGA (NC 13 to NC 14) in *WT H2Av* embryos vs Log2FC (p<0.05) between *Double H2Av* compared to *WT H2Av.* (A) Definition of quadrants QI-QIV. (B) Scatter plot of DEGs at NC 1. (C) Scatter plot of DEGs at NC 13. (D) Scatter plot of DEGs at NC 14. Numbers denote number of transcriptional units in each quadrant (QI– QIV). Panels B, C, and D also show the correlation coefficient (r) and p-value for the Pearson’s correlation test.

Of the 1496 transcriptional units significantly differentially expressed in NC 13, a larger portion were in quadrants QI and QIII (1394) than were in quadrants QII and QIV (102) (Figure 3C). QI transcriptional units are those that are developmentally upregulated and further upregulated in *Double H2A*v embryos, consistent with expedited or enhanced transcription. QIII transcriptional units are those that are developmentally downregulated and further downregulated in *Double H2Av* embryos, possibly due to accelerated transcript degradation. Thus, increased *H2Av* dosage shifts the transcriptional program of 94% of differentially expressed transcriptional units towards a more expedited developmental state. In contrast, only 6% of such transcriptional units displayed a pattern consistent with delayed development. Consistent with these findings, the Log2FC in *Double H2Av* vs *WT* embryos was positively correlated with the developmental change in WT (correlation coefficient r =∼0.70, p-value < 2.2e-16).

For NC 14, a comparable pattern emerged, though less pronounced (Figure 3D): 83% of the 1251 differentially expressed transcriptional units fell into QI and QIII, with a significant positive correlation between Log2FC in *Double H2Av* and developmental change (r = 0.2613925). For NC 1 only 176 were differentially expressed (Figure 3B), again consistent with the notion that the differences between the transcriptomes of the two genotypes largely develop as embryogenesis proceeds. Together, our data suggest that H2Av modulates the temporal regulation of gene expression during ZGA. Furthermore, QIII transcriptional units in Figure 3C cluster near the diagonal in the comparison plot, indicating that expression levels in NC 13 *Double H2Av* embryos already approximate those reached in *WT H2Av* embryos only at NC 14.

To identify the biological processes affected by this altered gene expression, we performed Gene Ontology (GO) enrichment analysis using Metascape. Precociously upregulated and downregulated transcriptional units (QI and QIII) were significantly enriched for fundamental cellular processes, including cell cycle regulation and RNA metabolism. In contrast, QII and QIV transcriptional units were enriched for developmental processes such as neurogenesis and tissue morphogenesis (Figure S5). Together, these results indicate that elevated *H2Av* dosage preferentially expedites the timing of housekeeping gene expression programs during ZGA, while largely leaving downstream developmental pathways unaffected.

Regarding the timing of mitosis 13, reducing and increasing levels of H2Av had opposite effects (Figure 2B). To determine if lower H2Av levels similarly change the timing of transcriptome remodeling, we performed a second round of RNA-Seq experiments. To be able to directly compare the results from these experiments with additional conditions (to be described below), we selected embryos of four genotypes (*Double H2Av*, *WT H2Av, Half H2Av*, *Half Jabba*) from three timepoints (NC 1, NC 11, NC 13) and performed RNA-Seq analysis on single embryos, as above. In the next paragraph, we will discuss the NC 1 and NC 13 comparison for *Double H2Av*, *WT H2Av,* and *Half H2Av* – other comparisons will be discussed in subsequent sections.

Analyzing this second set of RNA-Seq data, we categorized genes based on their canonical behavior during the MZT as either ZGA-upregulated (activated during zygotic genome activation) or ZGA-downregulated (targets of maternal RNA clearance) and computed the Log2FC in expression levels relative to *WT H2Av* at the same timepoint (Figure 4). For *Double H2Av* embryos, the pattern at NC 13 was similar to what we observed in our prior round of RNA-seq (Figure 3): ZGA-upregulated genes were on average further upregulated and ZGA-downregulated genes were further downregulated, indicating expedited transcriptome remodeling. *Half H2Av* embryos at NC 13 displayed the opposite pattern: transcriptional units normally downregulated at ZGA remained elevated compared to *WT H2Av*, while those that are normally upregulated were lower than in the wild type. Thus, similar to mitosis 13 timing, lower H2Av levels seemed to delay transcriptome remodeling.

**Figure 4.**
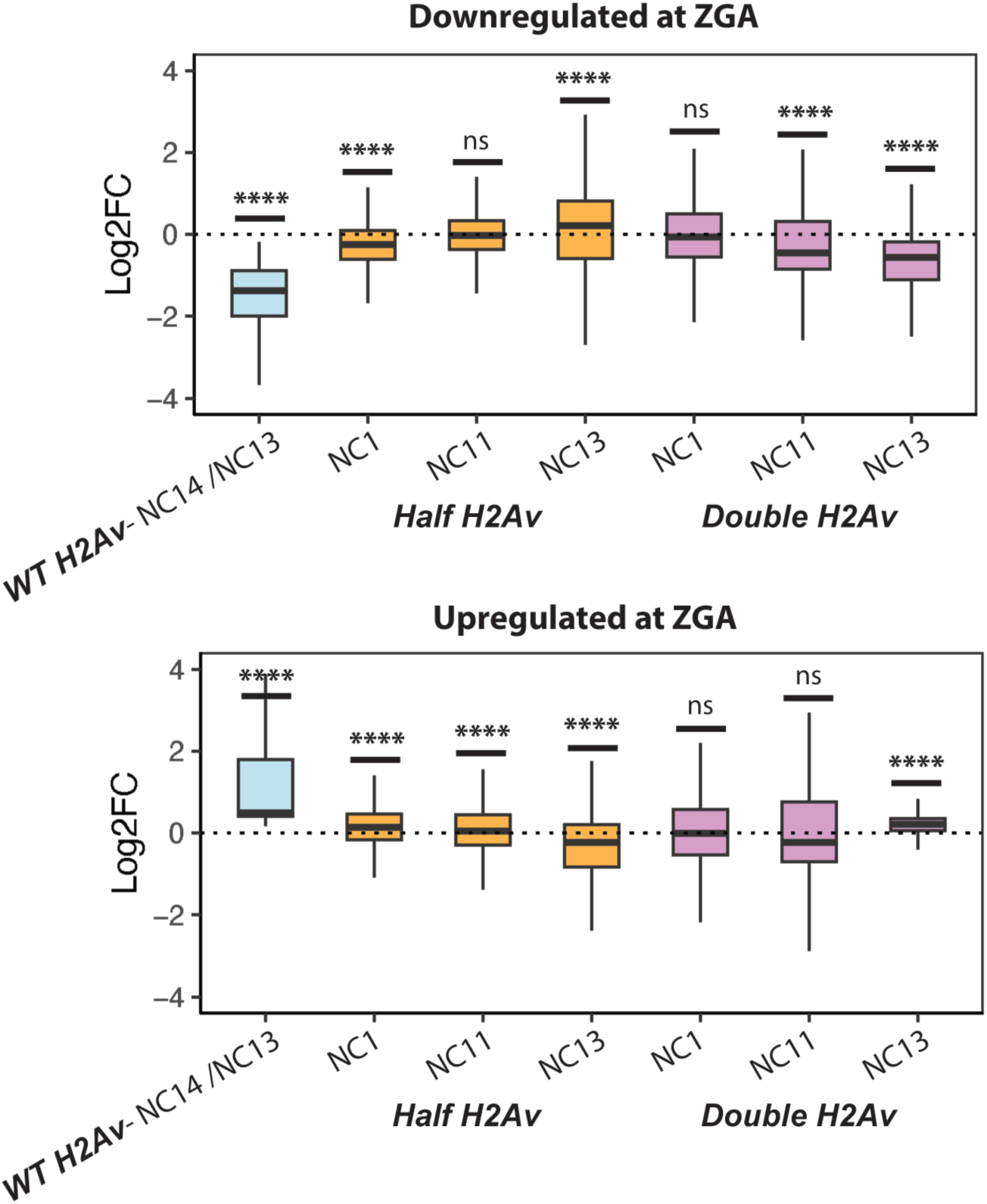
*H2Av* dosage modulates the timing of maternal transcript turnover and zygotic transcriptional activation. Box plots showing log₂ fold-change distributions for transcriptional units that are (top) ZGA-downregulated (normally net degraded during NC 13-NC 14) and (bottom) ZGA-upregulated (normally net increased during NC 13-NC 14) shown with blue boxes, comparing *Half H2Av* (orange boxes) and *Double H2Av* (pink boxes) embryos to *WT H2Av* controls at the indicated stages. Boxes indicate interquartile range and horizontal lines mark medians. Statistical significance was determined using the Wilcoxon/Mann–Whitney test. * for P ≤ 0.05, ** for P ≤ 0.01, *** for P ≤ 0.001, and **** for P ≤ 0.0001, “ns” = not significant (P ≥ 0.05). n = 3 independent biological replicates.

### H2Av-Dependent expression changes precede major zygotic genome activation

The eggs of *WT H2Av*, *Double H2Av*, and *Half H2Av* embryos are endowed with very similar levels of maternal mRNAs but their transcriptomes diverge dramatically by the ZGA (Figure 4). To characterize how these differences arise, we examined the transcriptomes of these genotypes at NC 11, which precedes the major transcriptional burst of the MZT. As above, we analyzed the expression levels of ZGA-upregulated and ZGA-downregulated genes relative to wild type. For *Double H2Av* embryos, transcriptional units normally downregulated exhibited a pronounced reduction in abundance relative to *WT H2Av* embryos (median Log2Fold change [Log2FC] = –0.45). This decrease was broadly observed across transcriptional units targeted for degradation at the MZT, indicating expedited maternal mRNA turnover prior to ZGA. Intriguingly, ZGA-upregulated genes were not significantly changed. This is consistent with the possibility that replenishment of these transcriptional units by zygotic transcription is modest at this timepoint and is outweighed by an accelerated turnover due to the increased H2Av levels. If so, increased H2Av levels may promote expedited mRNA degradation of an even broader range of transcriptional units, an effect partially masked at the NC 13 timepoint because of enhanced zygotic replacement. Again, *Half H2Av* embryos exhibited reciprocal effects. ZGA-upregulated transcriptional units were modestly elevated at NC 11 (median Log2FC = +0.05), whereas ZGA-downregulated genes showed little or no change relative to WT. These data indicate that a reduced dosage of *H2Av* delays maternal mRNA clearance.

Together, these observations support a model in which H2Av regulates early transcript dynamics in a dosage-dependent manner. Elevated H2Av enhances cytoplasmic maternal mRNA degradation prior to ZGA, whereas reduced H2Av slightly delays transcript turnover. The differential effects on ZGA-upregulated and ZGA-downregulated genes suggest that H2Av acts primarily through RNA stability rather than direct transcriptional repression at these stages.

### The nuclear pool of H2Av is not the main driver of transcriptome remodeling at the ZGA

We next investigated how H2Av levels control transcriptome remodeling. During NC 1 to NC 14, both global and nuclear H2Av levels rise (Figure 1). As H2Av is critical for the zygotic transcription of many genes (23), a straightforward model is therefore that zygotic transcription of certain genes depends on the levels of H2Av in the nucleus; these genes in turn control the rate of maternal mRNA decay and the timing of NC 13. However, H2Av is also abundant in the cytoplasm, stored on lipid droplets (18, 41); in principle, this non-nuclear pool could regulate the timing of embryonic processes via a non-chromatin mechanism. For histone H3, it has indeed been shown that its levels control cell-cycle progression around the MZT via a non-chromatin mechanism, by acting as a competitive inhibitor of the cell cycle kinase Chk1 (16). Since changing *H2Av* dosage alters both global H2Av levels and the nuclear pool in the same direction, it is impossible for us to distinguish if the effects of *H2Av* dosage on embryonic timing are mediated by either the increase in the total or the nuclear pool of H2Av. We therefore used two different strategies to determine if altering the nuclear pool of H2Av is sufficient to affect the timing of the cell cycle or transcriptome remodeling.

In our first approach, we took advantage of mutants of the histone chaperone Jabba which sequesters H2Av (as well as H2A and H2B) in the cytoplasm by recruiting it to lipid droplets. During oogenesis, this sequestration protects H2Av from degradation (18, 42); and during embryogenesis, it slows the import of H2Av into nuclei (17, 20). In the complete absence of Jabba, global H2Av levels in early embryos are therefore dramatically reduced while nuclear levels in NC 11-NC 13 are increased; H2Av associated with chromatin is also increased, as assessed by imaging of mitotic chromosomes (20). Intermediate effects are observed in embryos from mothers with reduced *Jabba* dosage (41, 42), in the following called *Half Jabba* embryos. Because these *Half Jabba* phenotypes had previously only been analyzed at a few timepoints, we measured global and nuclear H2Av levels in these embryos at the same timepoints as for *Half H2Av*, *WT H2Av*, and *Double H2Av*. Live imaging of embryos expressing H2Av–Dendra2 revealed an increase in nuclear H2Av signal in *Half Jabba* embryos during NC 13 compared to *WT H2Av* (Figure 5A,B), approaching those in *Double H2Av.* Western blot analysis across developmental stages showed that total H2Av protein levels were reduced relative to *WT H2Av* (Figure 5C,D). These findings indicate that although total H2Av abundance is decreased in *Half Jabba* embryos, a greater proportion of the available H2Av pool is incorporated into nuclei. Using this genotype, one can therefore interrogate if developmental timing is driven specifically by the nuclear pool of H2Av.

**Figure 5.**
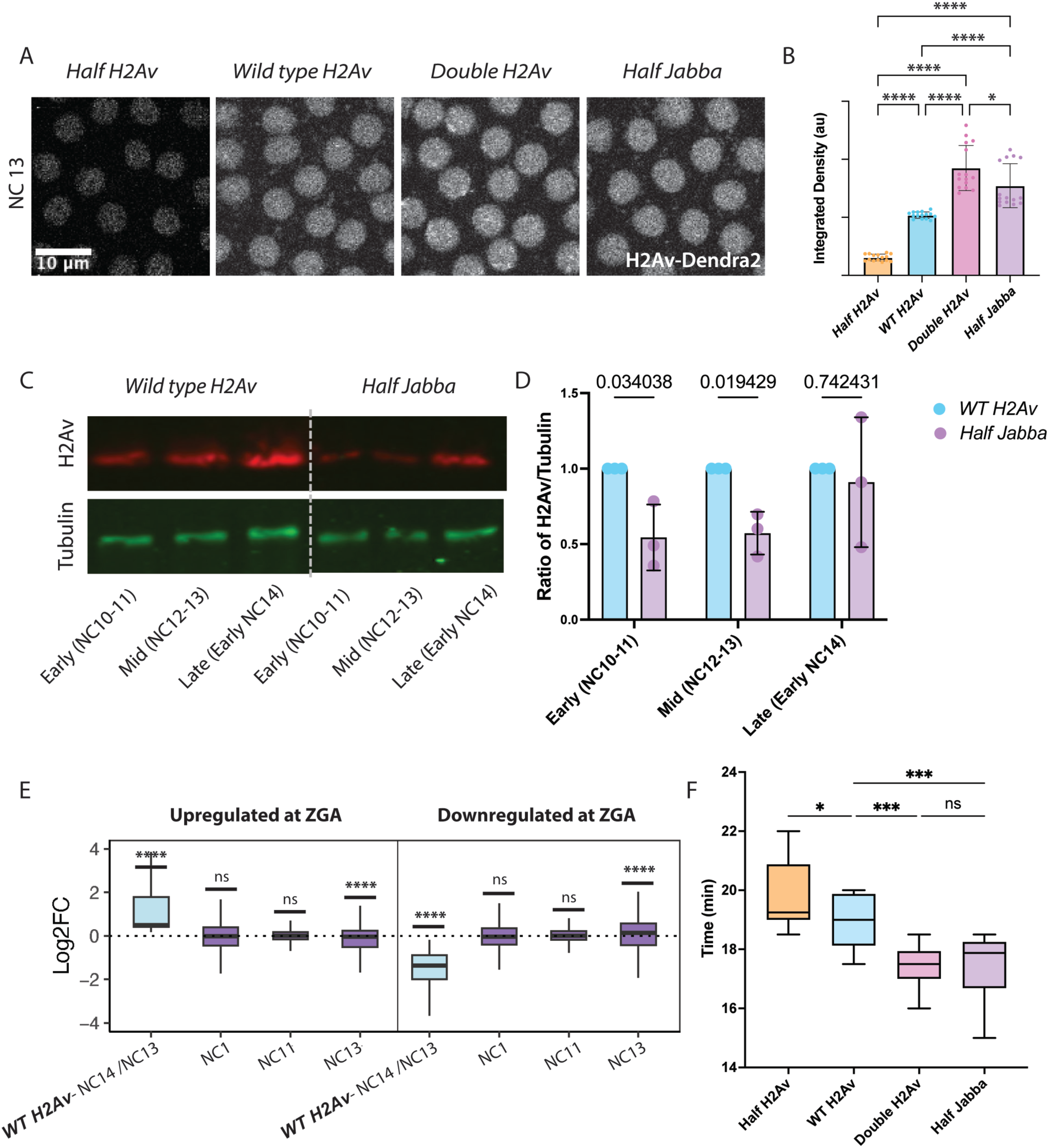
Altered H2Av distribution via decreased Jabba dosage speeds up NC 13 and delays transcriptional landscape. (A) Representative z-projections of Drosophila embryos at NC 13 expressing H2Av-Dendra2, illustrating nuclear H2Av accumulation. Scale bar: 10µm (B) Quantitation of sum of nuclear H2Av-Dendra2 fluorescence intensity per nucleus calculated at interphase of NC 11-NC 14. 3 nuclei were measured per embryo, N=3 embryos. Statistical analysis was performed using ordinary one-way ANOVA followed by Tukey’s multiple comparisons test. (C) Total H2Av protein levels are reduced in *Half Jabba* embryos compared to *WT H2Av* at NC 10-NC 13. The same Western blots were analyzed as in Figure 1, but just *WT H2Av* and *Half Jabba* are shown for clarity. Protein extracts from 10 visually staged embryos per condition (*Half Jabba* and *WT H2Av*) at Early (NC 10-NC 11), Mid (NC 12-NC 13), and Late (early NC 14) blastoderm were analyzed by SDS-PAGE and Western blotting. Membranes were probed with antibodies against H2Av (red) as well as against alpha-tubulin (green), as a loading control. (D) Quantitation of Western blots like in (C) for *Half H2Av*, *WT H2Av*, and *Double H2Av* embryos normalized to *WT H2Av* signal at each corresponding stage. Error bars represent standard deviation (n=3). Statistical analysis was performed using two-way ANOVA followed by Tukey’s multiple comparisons test. (E) Box plot of genes upregulated during the ZGA (left) and downregulated during ZGA (right). Line at median. Statistical analysis was performed using Wilcoxon/ Mann-Whitney test. ****=p<.0001. N=3 embryos. (F) Box plot showing NC timing in *WT H2Av* and *Half Jabba* embryos revealing faster progression through NC 13 in *Half Jab*ba embryos. Data for *Half H2Av*, *WT H2Av,* and *Double H2Av* embryos are the same as Figure 2B but shown again in this context for comparison to *Half Jabba*. Line at median. P values were calculated using ordinary one-way ANOVA followed by uncorrected Fisher’s LSD. * for P ≤ 0.05, ** for P ≤ 0.01, *** for P ≤ 0.001, and **** for P ≤ 0.0001, “ns” = not significant (P ≥ 0.05). N=10-12 embryos

We first measured progression through the syncytial nuclear cycles in *Half Jabba* embryos by live imaging. Mitosis 13 was expedited, similar to *Double H2Av* embryos (Figure 5F). Thus, the timing of NC 13 is likely driven by nuclear H2Av levels. To characterize the transcriptome of *Half Jabba* embryos, we sampled embryos from three timepoints (NC 1, NC 11, and NC 13) for *WT H2Av* and *Half Jabba* and performed RNA-Seq. Figure S8A compares the developmental trajectory of DEGs between *Half Jabba* and *WT H2Av* at NC 13. Recall that for *Double H2Av*, the behavior of 93% (1394 out of 1496) of the significantly changed transcriptional units was expedited (Figure 3). However, for *Half Jabba*, fewer than half (44%) were expedited, with the majority showing delayed behavior. Not surprisingly, there was also no significant correlation between the Log2FC of *Half Jabba* vs *WT* embryos and the developmental change in WT. This disparate pattern also held when we compared transcript levels overall: Figure 5E shows the levels of transcriptional units in *Half Jabba* embryos relative to *WT H2Av* embryos at three developmental stages, separately for transcriptional units upregulated or downregulated at ZGA. Even though nuclear H2Av levels of *Half Jabba* embryos approach those of *Double H2Av* embryos, the transcriptomes of the two genotypes showed very different patterns. For genes normally upregulated during ZGA, *Half Jabba* embryos exhibited reduced expression (median Log2FC = –0.05; Figure 5E). This pattern is opposite to that seen in *Double H2Av* embryos, where increased *H2Av* dosage promotes early activation. Transcriptional units normally downregulated during ZGA were upregulated in *Half Jabba* embryos at NC 13 (median Log2FC = +0.14; Figure 5E), even though they are further downregulated in *Double H2Av* embryos. Finally, when we determined if – compared to *WT H2Av* – transcriptional units in *Half Jabba* and *Double H2Av* changed in the same direction, we again observed a mixed pattern: out of 1752 transcriptional units, 56% showed concordant (QI and QIII) and 46% showed discordant (QII and QIV) behavior (Figure S8B,C). Together, these data indicate that nuclear H2Av levels can at most explain only a portion of the accelerated transcriptome remodeling observed in *Double H2Av* embryos.

In contrast, the transcriptomic pattern of *Half Jabba* embryos resembled more closely that of *Half H2Av* embryos. In both genotypes, downregulated transcriptional units tended to remain elevated compared to *WT H2Av*, corresponding to delay in transcriptome remodeling. The same is true for upregulated transcriptional units: for both genotypes, levels lagged behind those observed in the *WT H2Av*. Together, the pattern of transcript remodeling in *Half Jabba* embryos argues that it is the global, rather than the nuclear H2Av levels that mediate the timing effect on the embryonic transcriptome. We therefore speculate that the levels of cytoplasmic H2Av affect the stability of a subset of mRNAs, with increased levels in *Double H2Av* embryos promoting mRNA decay and reduced levels (likely for *Half Jabba* and *Half H2Av*) delaying decay.

As an orthogonal approach for analyzing the effects of nuclear H2Av on the transcriptome, we took advantage of a previous study which had knocked down the levels of the H2Av chaperone Domino during oogenesis (23). In these *Domino knockdown* (*DominoKD)* embryos, nuclear H2Av levels are markedly depleted. Nevertheless, analysis of steady-state RNA-seq profiles at NC 14 had revealed only modest changes to transcript levels of most genes (Figure S6A-C). Indeed, when we reanalyzed their data using the genes we have defined at either up– or down-regulated at ZGA, we found a median reduction in levels of only around –0.07 (Figure S6C). In addition, when we plotted the differences between the *DominoKD* and wild-type embryos against the transcript changes during normal ZGA, we found that most of the 6124 transcriptional units detected in both their and our dataset displayed less than two-fold changes in level (Figure S6A). When we applied a threshold of padj < 0.05, only ∼11% of transcriptional units (690) were significantly altered, with 392 downregulated and 298 upregulated (Figure S6B). Of these, the vast majority (593, 86%) fell into quadrants QI and QIII, suggesting that – if anything – their transcript remodeling is accelerated, similar to *Double H2Av* embryos, rather than delayed.

The same study investigating *H2Av* function performed global run-on sequencing (GRO-Seq) to specifically uncover effects of *Domino* knockdown on transcription. Here, the effects were much more profound. Transcription from many genes was significantly reduced (23). We reanalyzed these data focusing on the transcriptional units that are up– or downregulated at ZGA. Remarkably, we found massive downregulation in *DominoKD* embryos with more than 96% significantly down relative to *WT* (Figure S6D-F). These data indicate that many genes whose maternal transcripts change during the ZGA show substantial zygotic transcriptional contributions that depend on H2Av, consistent with earlier findings that numerous maternally deposited mRNAs possess zygotic transcriptional components (38–40).

Taken together, our re-analysis of the *DominoKD* data supports the idea that while lowering nuclear H2Av levels drastically affects the zygotic transcription of many genes it has only modest effects on steady-state RNA levels. Thus, both the *Jabba* and *Domino* data indicate that nuclear H2Av levels only have a modest effect on the global transcriptome, suggesting many of the dramatic transcriptome changes we observed in *Double H2Av* embryos arise by a non-nuclear mechanism.

### Increased H2A.Z accelerates the transcriptional landscape of zebrafish embryo

To test whether the role of H2A.Z in developmental timing is conserved, we examined the effect of *H2A.Z* dosage in a vertebrate, the zebrafish *Danio rerio*. We employed embryos with an additional transgenic copy of the H2A.Z coding sequence fused to GFP which is expressed from the endogenous *H2afz* promoter (43); this genotype is hereafter referred to as *H2A.Z-OE* (for over-expression). As zebrafish has two *H2A.Z* genes per haploid genome (44), the *H2A.Z-OE* line represents a 1.5-fold increase in *H2A.Z* dosage. We collected both WT and *H2A.Z-OE* embryos 2.5 hours and 4 hours post fertilization (hpf). These timepoints correspond to early and late MZT.

Similar to the strategies we employed for Drosophila, we plotted developmental changes (Late/Early MZT, WT) against genotype effects at early MZT (*H2A.Z-OE/WT*), generating four transcriptional quadrants (Figure 6A). A large portion of genes exhibited expedited upregulation in *H2A.Z-OE* embryos (QI; 1550 genes), indicating precocious activation of ZGA targets. As in our Drosophila studies, a substantial fraction of likely maternal transcripts exhibited enhanced downregulation (QIII; 1247 genes), consistent with accelerated clearance. QII (957 genes) and QIV (941 genes) contained smaller sets showing delayed degradation or reduced activation, respectively. Together, QI and QIII accounted for about 60% of affected transcriptional units, demonstrating that elevated H2A.Z expedites the transcriptional program of zebrafish embryos in a manner strikingly similar to Drosophila. Consistent with this notion, the genotype effects were significantly positively correlated with the expression level changes during development (r = 0.355467).

**Figure 6.**
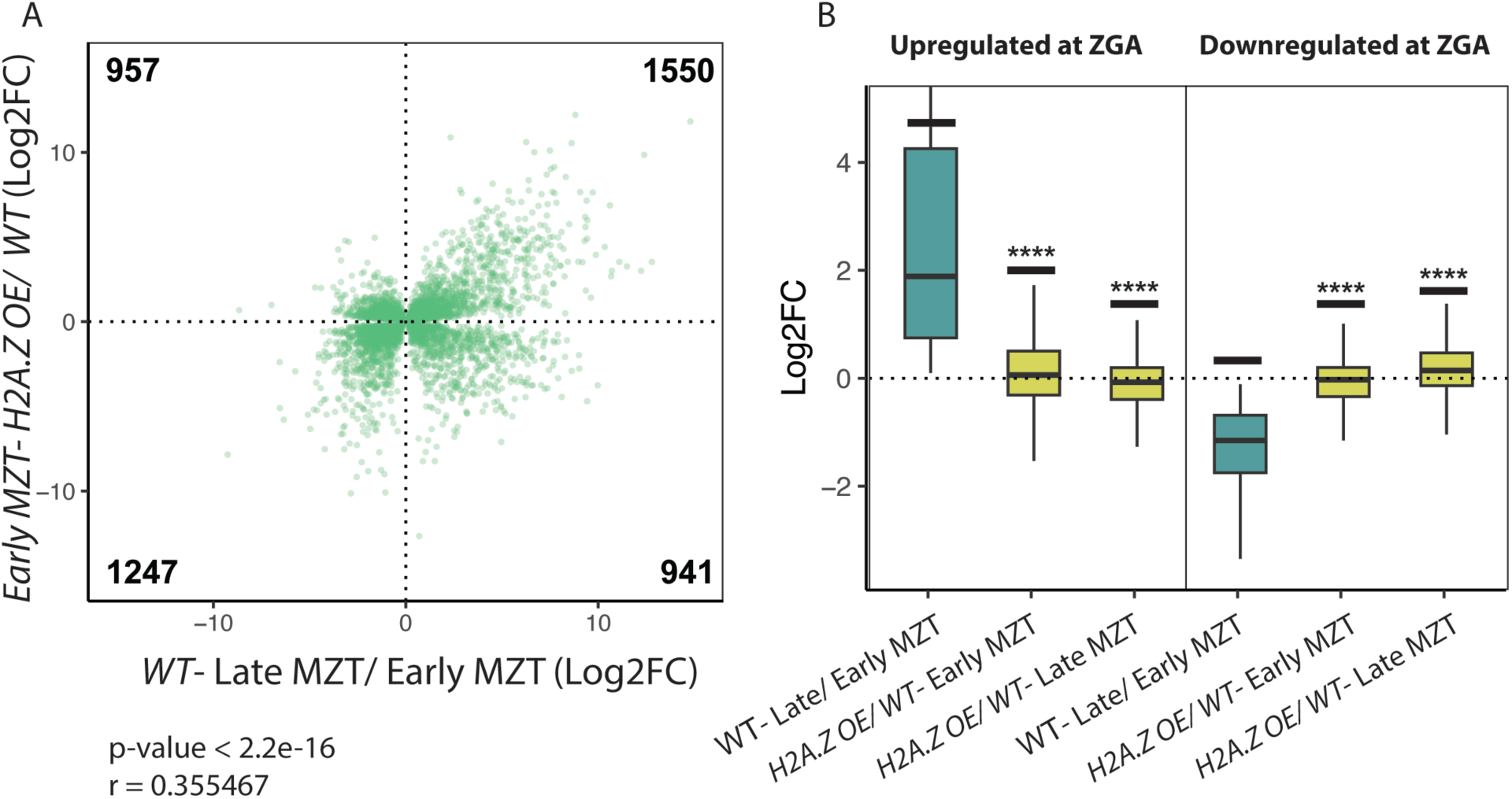
Overexpression of H2A.Z in zebrafish recapitulates the expedited transcriptome observed in *Drosophila*. (A) Transcriptional units were plotted based on their Log2FC from zero (p<0.05) early to late MZT in *WT H2A.Z* embryos vs Log2FC (p<0.05) between *H2A.Z OE* compared to *WT H2A.Z* at early MZT. (B) Box plots of genes upregulated during ZGA (left) and genes downregulated during ZGA (right), comparing H2A.Z-overexpressing and wild-type embryos. Lines at median. Statistical analysis performed using Wilcoxon/Mann–Whitney test. * for P ≤ 0.05, ** for P ≤ 0.01, *** for P ≤ 0.001, and **** for P ≤ 0.0001, “ns” = not significant (P ≥ 0.05). N = 3

Compared with *WT* early-MZT embryos, *H2A.Z OE* early-MZT embryos showed a modest increase in the Log2FC of transcriptional units that are normally upregulated during ZGA (median = 0.06). By late MZT, however, these same transcriptional units exhibited a slight decrease (median = –0.07). Transcriptional units normally downregulated during ZGA displayed the opposite pattern, early-MZT *H2A.Z-OE* embryos showed a small decrease (median = –0.02), whereas late-MZT embryos showed an increase (median = 0.14) (Figure 6B). These patterns indicate that H2A.Z accelerates both activation and clearance, producing an early boost in transcriptional dynamics that begins to normalize or slightly invert by late MZT. Thus, elevated H2A.Z primarily expedites the timing of the transcriptional program rather than permanently altering the magnitude of gene expression, a trend that mirrors observations in Drosophila embryos.

Thus, across both Drosophila and zebrafish developing embryos, increased *H2A.Z* dosage drives expedited accumulation of ZGA transcriptional units and expedited clearance of maternal RNAs, collectively accelerating the reprogramming in transcriptional landscapes that occur prior to major genome activation. Despite differences in embryonic architecture and MZT regulation, the direction of transcriptome shifts is conserved across these species. These findings identify H2A.Z as a shared temporal dial of early embryogenesis.

## Discussion

While the earliest stages of embryogenesis are driven by maternally provided factors, the control of later stages is dominated by transcription of the zygotic genome (1, 2). The transition between these stages, the MZT, involves many events occurring in parallel, such as cell cycle lengthening, degradation of maternal mRNAs, and large-scale ZGA, but it remains incompletely understood how these processes are temporally coordinated. Here, we find that in Drosophila expression levels of the variant histone H2Av time a large subset of events during the MZT, identifying H2Av as a tunable regulator of both nuclear cycle progression and transcriptome remodeling. During the MZT, both global and nuclear H2Av levels rise progressively. When this increase is experimentally accelerated via elevated *H2Av* dosage, the timing of mitosis 13 as well as the developmental trajectory of thousands of transcriptional units is expedited; a slower rise in H2Av levels leads to a corresponding delay in these events. We find that effects of *H2Av* dosage are mediated through distinct mechanisms: a nuclear H2A.Z–like activity that controls the timing of mitosis 13 and a non-nuclear role that dominates transcriptomic changes, likely via modulating maternal mRNA turnover. Finally, we provide evidence that this role as developmental timer is likely conserved: in zebrafish embryos, increasing the dosage of *H2A.Z* results in expedited increase or decrease of thousands of transcriptional units. We propose that levels of H2A.Z act as a conserved, critical regulator for the timing of many MZT events, providing a means for them to be executed in a coordinated fashion.

### Nuclear H2Av levels specifically modulate the timing of mitosis 13

Nuclear H2Av levels rise progressively from NC 11 to NC 14 (Figure 1C-D) (17, 20). Our data now reveal that this accumulation is not merely a passive consequence of development but actively contributes to the temporal regulation of cell cycle length, specifically in NC 13 (Figures S3A-B, S7). By uncoupling the H2A.X– and H2A.Z-like functions of H2Av, we show that phosphorylation at serine 137 and thus the canonical DNA damage response function of H2A.X are dispensable for NC 13 timing. Although DNA damage signaling clearly contributes to nuclear falling, loss of H2A.X function does not phenocopy the delayed mitosis observed in *Half H2Av* embryos. In contrast, increasing the *H2A.Z*-like dosage alone is sufficient to expedite NC 13. Together, these data argue that H2Av regulates cell cycle timing not via H2A.X-mediated checkpoint activation and instead implicate its H2A.Z-like chromatin function in setting the pace of embryonic divisions.

The mechanism(s) by which H2A.Z controls mitosis 13 timing remain to be elucidated. One possibility is that H2A.Z levels affect replication dynamics, *e.g.*, by altering the temporal pattern of when origins of replication are activated. In mouse stem cells, H2A.Z is enriched at origins (45); and in human cancer cells, it is critical for establishing early replication origins, as well as for controlling cell cycle length and proliferation (46). Alternatively, nuclear H2A.Z might regulate the zygotic transcription of a gene involved in promoting completion of NC 13. While globally the transcriptional units upregulated at ZGA are altered in opposite directions in *Half Jabba* and *Double H2Av* embryos (Figure 5E), a considerable subset changes in the same direction; *e.g.*, 132 transcriptional units are upregulated at least two-fold relative to wild type in both genotypes (Figure S8B). These transcriptional units are candidates for genes that speed up the progression of NC 13. Intriguingly, there are also 328 transcriptional units that are downregulated at least two-fold relative to wild type in both *Half Jabba* and *Double H2Av* embryos; these transcriptional units are enriched for cell-cycle genes (Figure S8B, C). Even though α amanitin injections to abolish zygotic transcription does not alter the duration of NC 13 (5, 30), it is conceivable that there are zygotic genes with opposing effect on NC 13 duration, some shortening and some delaying it. Indeed, embryos mutant for the pioneer transcription Zelda have widespread changes to their transcriptome (47, 48) and exhibit a delay in mitosis 13 (30). Thus, H2Av might selectively upregulate non-Zelda dependent zygotic genes that promote NC 13 completion.

### H2Av/H2A.Z levels regulate the pace of transcriptome remodeling in Drosophila and zebrafish

Despite major differences in embryonic architecture and regulatory logic between Drosophila and zebrafish, elevated H2Av/H2A.Z consistently promotes precocious activation of zygotic genes as well as accelerated clearance of maternal transcripts, suggesting a conserved role in the regulation of early embryogenesis. The seemingly more modest effect of *H2A.Z* dosage in zebrafish compared to Drosophila might simply reflect differences in experimental parameters: First, in our overexpression strains, *H2Av/H2A.Z* dosage is increased two-fold in Drosophila but only 1.5-fold in zebrafish. Second, the magnitude of the effect may depend on the timepoints analyzed. In Drosophila, the pattern for speed up is more obvious at NC 13 than at NC 14 (Figure 3C and 3D), presumably because earlier changes to the transcriptome lead to secondary effects at the later timepoint. Thus, examining additional stages of zebrafish MZT might result in more dramatic effects. Finally, our findings are consistent with our earlier studies on the H2A.Z chaperone Anp32e. In embryos lacking Anp32e, nuclear H2A.Z levels are increased, and many genes normally activated later in development are expressed precociously, consistent with early zygotic genome activation (36).

The dramatic effects of *H2Av/H2A.Z* dosage on the embryonic transcriptome have important technical implications. In Drosophila, fluorescently tagged H2Av is a convenient and widely used approach to monitor the nuclear cycle of early embryos live. Typically, this is done with tagged transgenes, thus increasing total *H2Av* dosage. Our findings argue that for early embryos transcriptomics data generated using such transgenes have to be interpreted with caution as the expression of hundreds of genes may be altered compared to strains with normal *H2Av* dosage; whether this distorting effect persists at other embryonic stages or even later in development remains to be determined.

### Drosophila H2Av levels regulate transcriptome remodeling in part via non-nuclear mechanisms

In Drosophila, *H2Av* dosage alterations only affect a subset of the transcriptome while the majority of transcriptional units is unaffected. Thus, the differences we observe are not due to a general change in the speed of development or errors when harvesting stages for analysis. Rather, H2Av levels specifically affect the levels of certain transcriptional units; these changes might result from changes in the new production of these transcripts or in their turnover.

Increased H2Av levels might stimulate or suppress the transcription of certain genes. Across organisms, H2A.Z is consistently enriched at promoters and poised enhancers, and thus well positioned to regulate transcription (32, 37, 49–51). In addition, H2Av can promote repression of genes, *e.g.*, through its role in heterochromatin formation (52–54). For transcriptional units typically upregulated during ZGA, higher *H2Av* dosage enhances their levels, consistent with transcriptional upregulation by H2Av. For transcriptional units typically downregulated, we observe the opposite effect, lower levels, consistent with more repression by H2Av. However, when nuclear H2Av levels are lowered by *Domino* knockdown, the transcription of both classes – as measured by GRO-seq – are reduced (23), arguing that H2Av is necessary to activate both. In addition, *Domino* knockdown has only relatively small effects on global RNA levels (23), suggesting that the bulk of changes we observe with altered *H2Av* dosage reflect changes to mRNA stability. We conclude that for the transcriptional units downregulated at ZGA H2Av promotes their turnover; this notion is consistent with the fact that we already observe effects at early timepoints when zygotic transcription is minimal. For the genes upregulated at ZGA, more H2Av might inhibit turnover and/or promote more transcription. Exactly how much each mechanism contributes will have to be addressed by future studies.

*How does H2Av promote the turnover of a subset of maternal transcriptional units?* A simple model is that H2Av is necessary for the transcription of zygotic genes that promote the degradation of these messages. For example, zygotic expression of microRNAs from the miR-309 cluster are known to promote the degradation of transcriptional units targeted by them, and in the absence of the micro RNAs, the turnover of these mRNAs is delayed (55). Premature expression of such microRNAs or other zygotic factors in *Double H2Av* embryos could lead to expeditated transcript clearance. However, three lines of evidence argue that such a model is not sufficient to explain the bulk of the changes we observe in mRNA turnover. First, *Double H2Av* embryos display lowered transcript levels already at NC 11, when zygotic transcription is limited. Second, total mRNA levels are only mildly affected in *DominoKD* embryos, even though lack of the hypothesized zygotic factor should reduce mRNA decay. Third, *Half Jabba* embryos have similar nuclear H2Av levels as *Double H2Av* embryos but have transcriptomic profiles resembling those of *Half H2Av* embryos rather than *Double H2Av* embryos. These findings indicate that nuclear H2Av is neither sufficient nor dominant in driving the accelerated maternal transcript clearance seen with increased *H2Av* dosage.

Our data point to a critical role of the total H2Av pool, rather than specifically the nuclear pool, in transcriptome remodeling during the MZT. Although nuclear H2Av levels in *Half Jabba* embryos approach those observed in *Double H2Av* embryos, their steady-state RNA profiles more closely resemble those of *Half H2Av* embryos: transcriptional units normally upregulated during ZGA show reduced activation, whereas transcriptional units normally downregulated remain elevated. We speculate that the cytoplasmic pool of H2Av present on lipid droplets somehow modulates the activity or availability of the RNA decay machinery, either directly or indirectly. Cytoplasmic histone stores are common in the early embryos of diverse species, including zebrafish (15, 35, 56) and Xenopus (14, 57), and thus might be able to influence early embryogenesis similarly via regulating maternal mRNA clearance. This hypothesis adds a new dimension to the models of MZT timing that have traditionally focused on titration of repressors of transcription (58–61) or RNA binding proteins (62, 63).

### A biological role for cytoplasmic H2Av sequestration

Newly laid Drosophila embryos store large amounts of histones H2A, H2B, and H2Av on lipid droplets, enough to package the DNA in over a thousand diploid nuclei (18). These histones are recruited to lipid droplets via Jabba (41), which protects them from proteasomal degradation during oogenesis (42) and buffers the import of H2Av into nuclei in early embryos (17, 20). The biological reason for this dramatic sequestration was previously obscure as *Jabba* mutant embryos develop grossly normally. These embryos take longer to hatch due to an unrelated function of Jabba in the spatial distribution of lipid droplets: late in NC 14, lipid droplets in *Jabba* mutants are misallocated to the yolk cell (64, 65); mutants with lipid-droplet misallocation but normal histone storage show a similar hatching delay. Other than that, *Jabba* mutants were only known to exhibit disruptions in the progression through early embryogenesis when exposed to additional stressors, such as reduced capacity to synthesize histones (41) or bacterial challenge (66). These observations might suggest that histone sequestration to lipid droplets mostly serves as a back-up system to enhance survival under unusual conditions.

The analysis described here shows that even under normal circumstances Jabba is needed for proper development. Reducing the capacity of histone storage in the *Half Jabba* embryos is sufficient to dramatically disrupt aspects of normal embryogenesis, including cell cycle progression and remodeling of the transcriptome. We conclude that Jabba-mediated sequestration of H2Av imposes a temporal delay on nuclear H2Av accumulation, providing a developmentally tunable mechanism to regulate the onset of zygotic transcription. The H2Av pool on lipid droplets may even regulate the mRNA decay machinery. While the mechanistic basis for these effects remains to be worked out, it is clear that Jabba has a dramatic effect on the embryonic transcriptome.

### H2Av as a tunable developmental timing dial

Taken together, our results support a model in which H2Av functions as a dosage-dependent dial that coordinates multiple aspects of early embryonic timing. Nuclear H2Av levels regulate the length of NC 13, while global H2Av abundance modulates the speed of transcriptome remodeling by influencing maternal mRNA stability. By acting through distinct nuclear and non-nuclear mechanisms, H2Av integrates chromatin state, cell cycle progression, and RNA dynamics to ensure orderly developmental transitions. Our work shows histone variant dosage as an underappreciated axis of developmental control and raises the possibility that modulation of histone pools more broadly contributes to temporal regulation during embryogenesis. Given that altered *H2A.Z* dosage also alters the timing of transcriptome remodeling in zebrafish, this function of H2Av/H2A.Z appears to be of ancient origin.

By tuning the exact levels of available H2A.Z in the nucleus and/or cytoplasm (*e.g.*, by how quickly *H2Av/H2A.Z* mRNA is translated or by sequestering H2Av/H2A.Z protein), embryos are in a position to regulate how quickly early embryogenesis proceeds. It will be an exciting prospect for future studies to test if such tuning of developmental speed is actually employed by embryos, either in response to environmental signals or due to evolutionary pressure. For example, if an organism were under evolutionary pressure to develop more rapidly, upregulating H2A.Z expression or tuning H2A.Z sequestration might provide an adaptive advantage.

## Materials and Methods

### Fly stocks

For westerns: Oregon R was used as *WT H2Av*, *w; H2AvKO/TMCB* (24, gift from Jürg Müller) as *Half H2Av*, and *gH2Av* (20) was used as *Double H2Av*. For imaging: Endogenously tagged *H2Av-Dendra2* was used as *WT H2Av*. To generate this line, endogenously tagged H2Av-Dendra2, a single CRISPR target site near the stop codon was selected using Target Finder (67) and the designed target gRNA sequence (GGTGCAGGATCCGCAGCGGA) was subcloned into the pU6-BbsI-chiRNA vector (a gift from Melissa Harrison, Kate O’Connor-Giles, and Jill Wildonger, Addgene plasmid #45946). A 1 kb left homology arm with a synonymous mutation at the PAM site and a 1 kb right homology arm were synthesized and assembled in the pScarless-Dendra2-3xP3-DsRed plasmid backbone (19) (GenScript). The plasmids of gRNA and homology arms were co-injected into nos-Cas9 embryos (TH00788.N) and DsRed+ progeny were screened (BestGene Inc., Chino Hills, CA). The DsRed marker was removed through a cross to nos-PBac flies (a gift from Robert Marmion and Stanislav Shvartsman) (68). Insertion of the fluorescent protein was confirmed by PCR and Sanger sequencing. The pMBAC-1xHisC plasmid was a gift from Robert J. Duronio; the pMBAC-1xHisC-H3-Dendra2 plasmid containing Dendra2 was previously described (19). Transgenic flies carrying a genomic *H2Av-Dendra2* construct on the second chromosome (20) were crossed *to Endo-H2Av-Dendra2* to produce flies expressing 4 copies of *H2Av-Dendra2* and used as *Double H2Av*. *Endo-H2Av-Dendra2* flies were crossed with w*; H2AvKO/TMCB* (24) to produce flies expressing 1 copy of *Endo-H2Av-Dendra2*, and used as *Half H2Av*. To generate *yw;sp/cyo;1X HisC H2A-Dendra2 (vk33),* Dendra2 was subcloned into the C-terminus of H2A to generate pMBAC-1xHisC-H2A-Dendra2 (Genscript). The transgene was inserted into the vk33 attP site via phiC31-mediated integration (BestGene Inc., Chino Hills, CA). *Jabba^DL^* (41) was used as a *Jabba null* allele. We used FlyBase to find information on genes, gene expression, stocks, and sequences (69).

### Microscopy/ live imaging

All imaging was performed on a Leica SP5 laser scanning confocal microscope with Leica HyD hybrid detectors. To measure nuclear H2Av-Dendra2 levels in embryos of various *H2Av* dosages, embryos were manually dechorionated and mounted on coverslips coated with a heptane-based glue (3M 667 Scotch® Double-Sided Tape dissolved in heptane), then overlaid with halocarbon oil 27 (Sigma-Aldrich, Burlington, MA). NC 10 embryos were selected and imaged until NC 14. Acquisition included 10 slices, 1µm apart, at 1 minute time intervals. Imaging was performed using a 63X oil immersion objective (NA 1.4) and 488nm excitation. To measure nuclear H2Av-Dendra2 levels in *Half Jabba* embryos compared to other genotypes, NC 13 embryos were imaged. Acquisition included 10 slices, 1µm apart.

### Nuclear cycle length measurement

NC 10 embryos were selected, mounted as above, and images were captured every 15 sec using a 40X oil immersion objective (NA 1.3). NC lengths were measured from one anaphase to the next anaphase.

### Nuclear intensity measurement

Quantification was performed by selecting a 32.23 x 32.23 µm region of interest (ROI) within the embryo, using a previously described method (19). Three consecutive slices with the greatest signal intensity were chosen for projection using the “Sum” function in ImageJ. Within the projected ROI, three individual nuclei were selected for analysis at NC 11 through 14.

### Immunofluorescence staining

Embryos were collected on apple juice agar plates for one hour, aged for an additional 2 hours, and dechorionated with 50% bleach. Embryos were then fixed in 4% formaldehyde in 1X PBS for 20 min, followed by devitellinization as described (70). NC 14 embryos were selected and blocked overnight at 4°C in a solution of 10% BSA, 0.1% Triton X-100, and 1X PBS. Primary antibody incubation was performed overnight at 4°C using anti-gammaH2Av antibody UNC93-5.2.1-s (1: 1000; Developmental Studies Hybridoma Bank, Iowa City, Iowa). Samples were washed in a solution of 1X PBS and 0.1% Triton X-100 and incubated with secondary antibody (1: 1000; goat anti-mouse Alexa 633 (Invitrogen, Carlsbad, California). Embryos were mounted in Aqua-Poly/Mount (Polysciences, Warrington, PA) using 18 mm X 18 mm coverslips as spacers.

### Nuclear falling quantification

Embryos aged about 2.5 hours post-fertilization were collected, dechorionated, heat fixed as described (71), and staged by nuclear cycle. NC 14 embryos were selected and blocked overnight at 4°C in a solution of 10% BSA, 0.1% Triton X-100, and 1X PBS. DNA was labeled using DAPI (Sigma, St. Louis, MO). Embryos were mounted and imaged as above using the 40x objective. Z-stack images were acquired, spanning 50 µm in depth at 1 µm intervals, from the embryo surface to interior above the yolk. To assess nuclear positioning, 3D0 renderings were generated in AMIRA (Thermo Fisher Scientific), and the number of internalized (fallen) nuclei was quantified along the y-z and x-y axes (72).

### Western blot analysis

Embryos were heat-fixed, staged, and boiled in Laemmli buffer (Bio-Rad, Hercules, CA) for 15 minutes. Protein samples were separated on 4%-15% SDS-PAGE gels (Bio-Rad) and transferred to PVDF membranes (Immobilon-FL, MilliporeSigma, Burlington, MA). Transfers were performed in Towbin buffer at 80V for 30 min. Membranes were probed with the following antibodies: mouse anti-H2Av (1:5000, Active Motif, Carlsbad, CA), rabbit anti-alpha-Tubulin (1:10,000; Cell Signaling, Danvers, MA). Corresponding secondary antibodies were: IRDye® 800CW goat anti-rabbit IgG, IRDye® 680RB, IRDye® 680RD donkey anti-guinea pig IgG, goat anti-mouse IgG (all 1:10,000; LI-COR, Lincoln, NE). Blots were imaged using a LI-COR Odyssey CLx system, and intensity was quantified using Image Studio Lite software.

### H2Av^S137A^-Dendra2 line generation

Site-directed mutagenesis was performed using the Agilent QuikChange II Site-Directed Mutagenesis Kit according to the manufacturer’s protocol. Complementary oligonucleotides carrying the desired mutation in *H2Av* (primer 1: 5’-ggc aac gtc att ctg gcg cag gcc tac taa gcc-3’; Primer 2: 5’-ggc tta gta ggc ctg cgc cag aat gac gtt gcc-3’) were used. A previously generated *H2Av-Dendra*2 transgene plasmid (20).was used as the dsDNA template. Reactions were transformed into XL10-Gold ultracompetent cells and plated on selective LB-agar. The insert was then amplified via PCR and ligated into the pattB plasmid. Transgenic lines were created using PhiC31 integrase-mediated transgenesis (BestGene Inc., Chino Hills, CA). All insertions were incorporated onto the third chromosome at site 68A4.

### Quantitation and image processing

All image analysis was performed using ImageJ. Statistics were performed using Prism7, GraphPad. All images are assembled with Adobe Illustrator.

### RNA-Sequencing

Drosophila: To clear fertilized but unlaid eggs from females, embryos were collected on plates with fresh yeast paste for one hour and discarded. Flies were then allowed to lay for 15-30 min, and collection plates were either immediately processed (NC 1) or aged at 25°C. Plates were covered with halocarbon oil 27 (Sigma-Aldrich, Burlington, MA) to turn the chorion transparent to allow for visual assessment of the embryonic stage. For the NC 1 timepoint, embryos exhibiting ubiquitous H2Av-Dendra signal from anterior to posterior (with the posterior membrane not yet retracted) were selected, and individual embryos were flash-frozen by placing them in an Eppendorf tube containing 1 µl of TRIzol (Invitrogen, Carlsbad, California) and then onto dry ice. For the other timepoints, embryos of roughly the correct stage were observed by confocal imaging until they reached the desired timepoint and then flash frozen. For the NC 11 timepoint, embryos were selected 5 min after the 10^th^ anaphase. For the NC 13 timepoint, embryos were selected at 8 min after the 12^th^ anaphase. For the NC 14 timepoint, embryos were selected at 15 min after the 13^th^ anaphase. Three embryos were collected for each experimental condition. Embryos were then macerated in TRIzol with a needle to extract RNA and purified using Direct-Zol™ RNA MiniPrep Kit (Zymo Research, Irvine, CA) according to manufacturer’s protocol. RNA sequencing was performed by GENEWIZ (South Plainfield, NJ). Libraries were prepared as Total RNA via Ribosomal RNA depletion.

Zebrafish: Wild-type and *H2A.Z-OE* embryos were harvested after 2.5 hpf and 4 hpf, dechorionated, and homogenized in TRIzol solution. RNA was precipitated from samples using phenol–chloroform and purified using Direct-Zol™ RNA MiniPrep Kit. RNA was submitted to University of Rochester Genomic Center for quality check and sequencing. In brief, RNA quality was evaluated using BioAnalyzer and samples with RNA integrity number > 9 were used for library synthesis. Libraries were synthesized using the Illumina Stranded Total RNA Prep with Ribo-Zero and sequenced with paired-end 150 bp reads on a NovaSeq X Plus platform.

### RNA-Sequencing analysis

There are four batches of RNA-seq data: Supplementary Table 1 summarizes RNA-seq samples and batch information. Generally, RNA-seq reads were trimmed for adaptors using TrimGalore (version 0.6.10) (73) and then mapped to the *D. melanogaster* reference genome (dm6) using STAR (version 2.7.11b). The transcript-level abundance was quantified with Salmon (74) and then aggregated using tximport (75) to obtain gene-level quantification. Subsequent differential expression analysis was done using DESeq2 (74). For batch four RNA-seq reads, we used SMARTer Stranded Total RNA Kit – Pico Input to prepare the libraries. Before adapter trimming, the UMIs in read2 of the paired-end RNA-seq were extracted using UMI-tools extract (version 0.5.5, ––bc-pattern=NNNNNNNNNNNNNN) (76). Then after adapter trimming and alignment, the duplicated reads were removed from the alignment files with UMI-tools dedup (version 0.5.5) (76). Then the alignment files were converted back to FASTQ format with SAMtools (version 1.20) (77) and used for downstream transcriptome analysis.

Batch information was used as covariant in differential expression analysis when different batches of RNA-seq data were combined. For *Domino* knockdown data from Ibarra-Morales, et al., 2021, the DESeq2 results were downloaded and plotted with our results. Statistical tests and plots for data visualization were generated in R and GraphPad Prism 10.

For the analysis of zebrafish RNA-seq data, raw sequencing reads were trimmed for adaptors using TrimGalore (version 0.6.10) (73), and then mapped to the zebrafish genome assembly (GRCz11) using STAR (version 2.7.11b) (78). The transcript-level abundance was quantified with Salmon (74) and then aggregated using tximport (75) to obtain gene-level quantification. Subsequent differential expression analysis was done using DESeq2 (79). Statistical tests and plots for data visualization were generated in R (73).

All RNA sequencing datasets generated in this study have been deposited to GEO data repository under accession number GSE327664 (reviewer access token: cfaligogdxkzbin).

### Gene ontology analysis

Gene ontology (GO) analysis was conducted using Metascape (80) with default settings. For QI, QII, QIII, and QIV maternal transcriptional units at NC 13, a single gene list was analyzed per group. Gene counts for each GO term were converted to gene ratios (observed count/ total genes in the term before visualization. The top 4-5 significant GO terms per group were selected and ordered by hierarchical clustering based on the distance of –log_10_p-values (80).

### Ethics Statement

Zebrafish care and maintenance were performed in accordance with IACUC approved animal care and use guidelines with ethical approval by the University Committee on Animal Resources at the University of Rochester Medical Center, and the Cornell Center for Animal Resources and Education.

## Acknowledgements

We thank Jürg Müller, Robert Duronio, and the Bloomington Drosophila Stock Center (NIH P40OD018537) for providing fly lines. We are grateful to Melissa Harrison, Kate O’Connor-Giles, Jill Wildonger, Robert Marmion, and Stanislav Shvartsman for reagents.

## Supplemental information

**Table S1.** RNA-seq summary.

**Figure S1:** H2A levels in nuclei decrease during early nuclear cycles while those of H2Av increase. (A, B) Developmental decrease of nuclear H2A. (A) Representative images of Drosophila embryos at nuclear cycles (NC) 11, NC 12, NC 13, and NC 14 expressing one tagged transgene of H2A-Dendra2, illustrating progressive reduction in nuclear accumulation of H2A. Scale bar: 10µm (B) Quantification of sum of nuclear H2A-Dendra2 fluorescence intensity per nucleus calculated at interphase of NC 11-NC 14. 3 nuclei were measured per embryo, N=3 embryos. Statistical analysis was performed using ordinary one-way ANOVA followed by Tukey’s multiple comparisons test. * for P ≤ 0.05, ** for P ≤ 0.01, *** for P ≤ 0.001, and **** for P ≤ 0.0001, “ns” = not significant (P ≥ 0.05). (C-E) Developmental increase of nuclear H2Av. Same data as in Figure 1, but here plotted to compare changes in nuclear H2Av levels from NC11-NC13 within each genotype. Quantitation of total nuclear H2Av-Dendra2 fluorescence intensity per nucleus during interphase of NC 11-NC 13 in (C) *Half H2Av*, (D) *WT H2Av*, and (E) *Double H2A*v embryos. Five nuclei were measured per embryo for three embryos per genotype. Error bars represent mean with SEM. Statistical analysis was performed using one-way ANOVA with Kruskal–Wallis test. * for P ≤ 0.05, ** for P ≤ 0.01, *** for P ≤ 0.001, and **** for P ≤ 0.0001, “ns” = not significant (P ≥ 0.05).

**Figure S2:** Analyzing nuclear falling in early embryos. (A) Experimental Design showing how nuclear falling was quantified. 50 µm Z-stacks of DAPI stained embryos were obtained (from the periphery of embryos to the middle of the embryo every 1 µm). 3D renderings were generated using AMIRA, and the number of internalized nuclei (NF) was counted in the y-z and x-y-planes. Boxes represent the view observed in cross section. White arrowhead= fallen. (B) Snapshot of live imaging of *Double H2Av-Dendra2* embryos showing a chromosome bridge (white arrowhead) between recently condensed nuclei. Scale bar= 10um. (C) DAPI (cyan) staining and gamma-H2Av (red) immunostaining of *Double H2Av* embryo reveal prominent gamma-H2Av staining in fallen nuclei, indicative of DNA damage. (D) Cross section of whole embryo. Scale bar represents 50um. Only nuclei extruded from periphery display appreciable gamma-H2Av signal. Scale bar= 10um.

**Figure S3:** H2Av has little effect on NC 11 and NC12 timing. Embryos of various genotypes were imaged at 25°C during NC 10-NC 14. The duration of nuclear cycles was quantified from time-lapse movies by measuring the interval between anaphase onset of one mitosis and the next. Box plots showing (A) NC11 and (B) NC12 duration in genotypes with varying *H2A.Z* and *H2A.X* dosage. Line at median. P values were calculated using ordinary one-way ANOVA followed by uncorrected Fisher’s LSD. * for P ≤ 0.05, ** for P ≤ 0.01, *** for P ≤ 0.001, and **** for P ≤ 0.0001, “ns” = not significant (P ≥ 0.05). N=11-12 embryos. (C) Schematic of a genomic transgene encoding non-phosphorylatable H2Av-Dendra2. The conserved serine at position 137 (equivalent to the DNA damage responsive phosphorylation site in H2A.X) is mutated to alanine (S137A), enabling separation of H2A.Z-specific functions from those of H2A.X.

**Figure S4:** Expression of many maternal and maternal-zygotic transcriptional units are altered in *Double H2Av* embryos. (A) Schematic of RNA-Sequencing pipeline, created using BioRender. Single embryos were visually staged via live imaging, flash frozen, and macerated in TRIzol. Total RNA was extracted and purified for transcriptomic analysis by RNA-Sequencing. (B-C’) Volcano plots showing differential gene expression in *WT H2Av* embryos between time points on x-axis. Significantly changing transcriptional units (p< 0.05) are colored by classification: maternal (red) and zygotic (blue). Non-significant transcriptional units are shown in grey and excluded from DEG counts (displayed on plots). Maternal and zygotic transcripts were defined based on transcript abundance (TPM) at NC 1: transcriptional units with TPM > 1 were classified as maternal, those with TPM = 0 as zygotic, and transcriptional units with 0 < TPM < 1 were excluded from analysis. Transcriptional units that are upregulated between the NC 1-NC 13 and NC 13-NC 14 timepoints are defined as tentative maternal-zygotic transcripts as their expression must be supplemented by embryonic transcription. (B) Differentially expressed maternal transcriptional units in *WT H2Av* embryos at NC 1-NC 13. Significantly downregulated=874; significantly upregulated= 791. (C) Differentially expressed maternal transcriptional units in *WT H2Av* embryos at NC 13-NC 14. Significantly downregulated=1931; significantly upregulated= 2265. (C’) Differentially expressed zygotic transcriptional units in *WT H2Av* embryos at NC 13-NC 14. Significantly downregulated=0; significantly upregulated= 555. (D-F’) Pie charts representing DEGs (p<0.05) in *Double H2Av* compared to *WT H2Av* at each time point (NC 1, NC 13, and NC 14). Percentages are of total detected maternal (6661) and zygotic (4395) transcriptional units. (D) 4.52% (301) maternal transcriptional units were differentially expressed at NC 1. (E) 27.77% (1850) maternal transcriptional units were differentially expressed at NC 13. (E’) 1.80% (79) zygotic transcriptional units were differentially expressed at NC 13. (F) 25.49% (1698) maternal transcriptional units were differentially expressed at NC 14. (F’) 1.91% (84) zygotic transcriptional units were differentially expressed at NC 14. (G) Differentially expressed maternal transcriptional units at NC 1. Significantly downregulated=137; significantly upregulated= 164. (H) Differentially expressed maternal transcriptional units at NC 13. Significantly downregulated=1141; significantly upregulated= 709. (H’) Differentially expressed zygotic transcriptional units at NC 13. Significantly downregulated=35; significantly upregulated= 44. (I) Differentially expressed transcriptional units at NC 14. Significantly downregulated=1173; significantly upregulated= 525. (I’) Differentially expressed zygotic transcriptional units at NC 14. Significantly downregulated=54; significantly upregulated= 30.

**Figure S5:** Quadrants I and III are enriched in housekeeping genes. Gene set enrichment analysis using Metascape showing top 5 most significant GO terms per quadrant (quadrants as defined in Fig. 3). X-axis measures log10 q-value and the size of the circle is proportional to the number of observed genes presented.

**Figure S6:** *Domino* knockdown causes modest steady-state RNA changes but strong transcriptional disruption at ZGA. (A-C) Analysis of RNA-Seq data from Ibarra-Morales et al., 2021. (A) Scatter plot of log2 fold change of gene expression for *DominoKD* compared to *WT* control for genes during ZGA (NC13-NC14). (B) Scatter plot in (A) with padj<0.05 thresholding. (C) Boxplot of log2 fold changes of DEGs in *Double H2Av* embryos that are upregulated (left) or downregulated (right) at ZGA (NC13-NC14) in *WT H2Av* embryos. Genes represented are those identified in Ibarra-Morales et al., 2021. Lines at median. Statistical analysis performed using Wilcoxon/Mann–Whitney test. (D-F) Analysis of GRO-Seq data from Ibarra-Morales et al., 2021. (D) Scatter plot of log2 fold change of gene expression for *DominoKD* compared to *WT* control for genes during ZGA (NC13-NC14). (E) Scatter plot in (D) with padj<0.05 thresholding. (F) Boxplot of log2 fold changes of DEGs in *Double H2*Av embryos that are upregulated (left) or downregulated (right) at ZGA (NC13-NC14) in *WT H2Av* embryos. Genes represented are those identified in Ibarra-Morales et al., 2021. Lines at median. Statistical analysis performed using Wilcoxon/Mann–Whitney test.

**Figure S7:** NC 11 and NC12 timing in *Half Jabba* embryos. *Half Jabba* embryos were imaged at 25°C during NC 10-NC 14. The duration of nuclear cycles was quantified from time-lapse movies by measuring the interval between anaphase onset of one mitosis and the next. Box plots showing (A) NC11 and (B) NC12 duration in *Half Jabba* embryos compared to other genotypes, revealing faster progression through NCs. Data for *Half H2Av*, *WT H2Av,* and *Double H2Av* embryos are the same as Figure S2 but shown again in this context for comparison to *Half Jabba*. Line at median. p values were calculated using ordinary one-way ANOVA followed by uncorrected Fisher’s LSD. * for P ≤ 0.05, ** for P ≤ 0.01, *** for P ≤ 0.001, and **** for P ≤ 0.0001, “ns” = not significant (P ≥ 0.05). N=10-12 embryos.

**Figure S8:** *Half Jabba* embryos exhibit both expedited and delayed transcriptome remodeling. (A) Comparison of DEGs in *Half Jabba* compared to *WT H2Av* at NC13 (padj<0.05). Significantly altered transcriptional units fall into all four quadrants; 44.2% (QI and QIII) display expedited and 55.8% (QII and QIV) display delayed behavior. The Pearson’s correlation test revealed no significant correlation. (B) Comparison of Log2FCs (relative to wild type) between *Half Jabba* and *Double H2Av* at NC13 (padj<0.05) for those transcriptional units that are either up– or downregulated at ZGA. (C) The same data in as in B, with candidate cell cycle genes highlighted. Blue dots in Q3 are transcriptional units identified as related to the regulation of cell cycle by GO analysis.

**Figure S9:** *Double H2Av embryos* have increased levels of chromatin-associated H2Av. Quantitation of chromatin-associated H2Av-Dendra2 fluorescence intensity per nucleus during metaphase of NC 11-NC 13 in *Half H2Av*, *WT H2Av*, and *Double H2Av* embryos. Five metaphase nuclei were measured per embryo for three embryos per genotype. Error bars represent mean with SEM. Statistical analysis was performed using two-way ANOVA followed by Tukey’s multiple comparisons test. * for P ≤ 0.05, ** for P ≤ 0.01, *** for P ≤ 0.001, and **** for P ≤ 0.0001, “ns” = not significant (P ≥ 0.05).

