## Supplemental materials for "H2A.Z levels control the timing of major events at the maternal-zygotic transition"

**Figure S1: H2A levels in nuclei decrease during early nuclear cycles while those of H2Av increase**

Figure S1

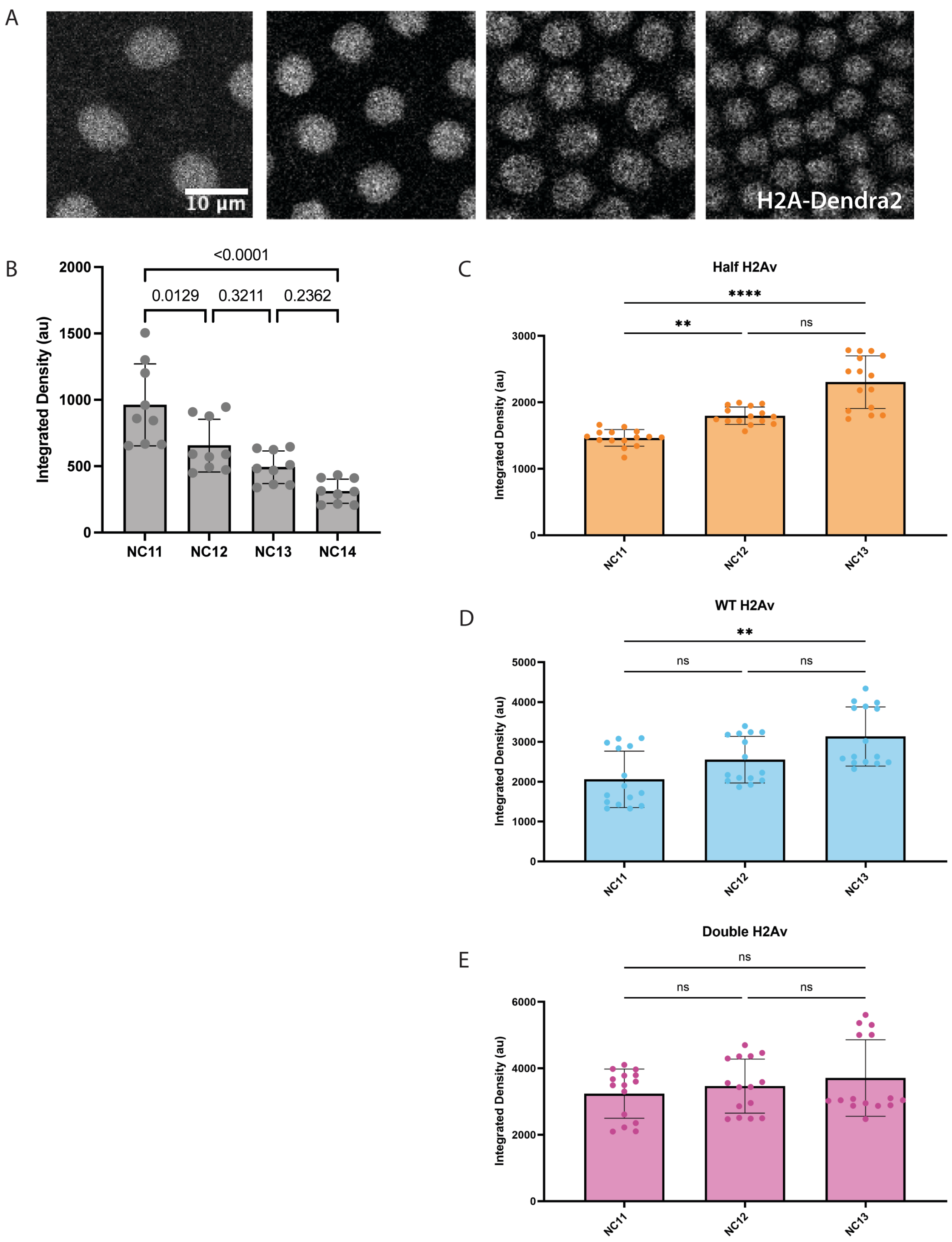

**Figure S2: Analyzing nuclear falling in early embryos.**

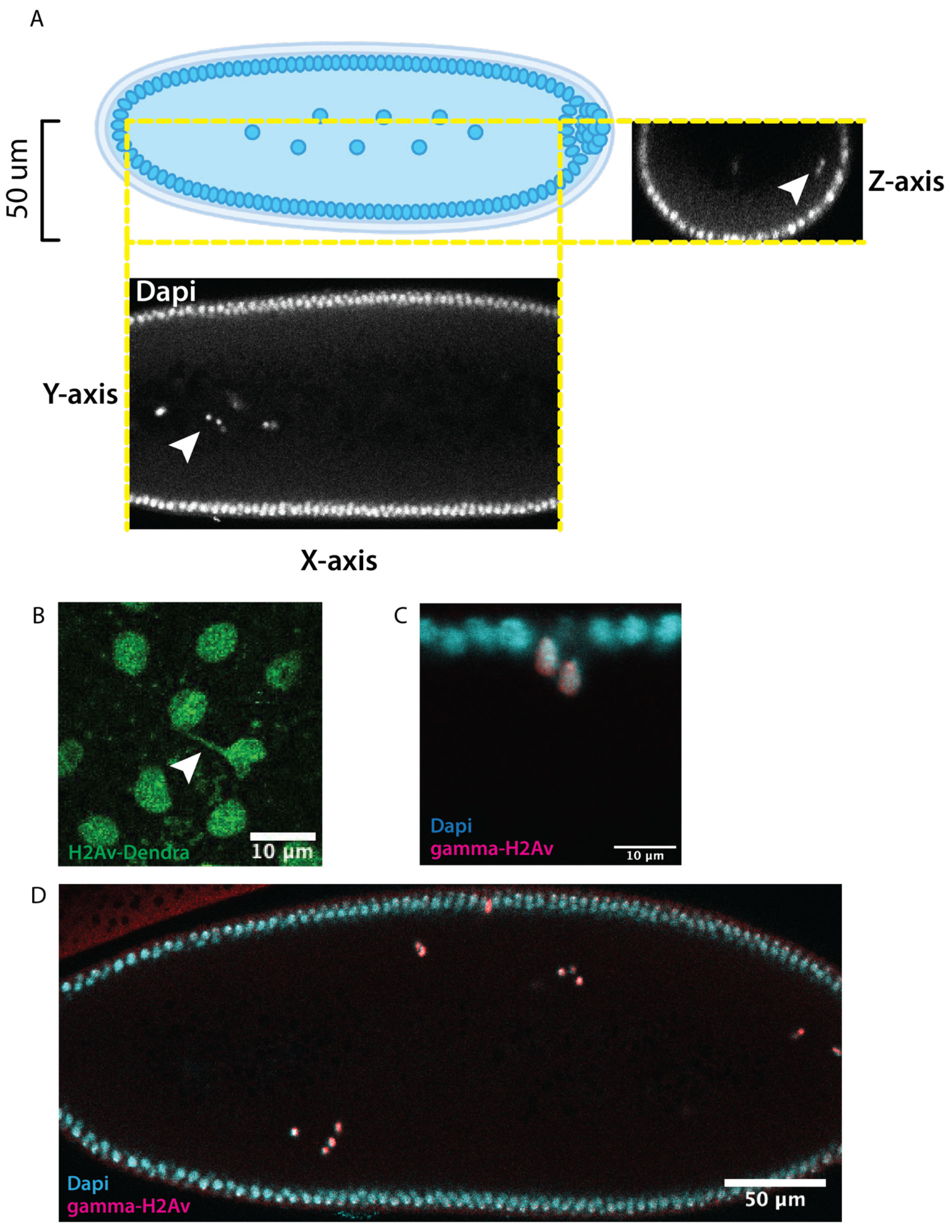

**Figure S3: H2Av has little effect on NC 11 and NC12 timing**

Figure S3

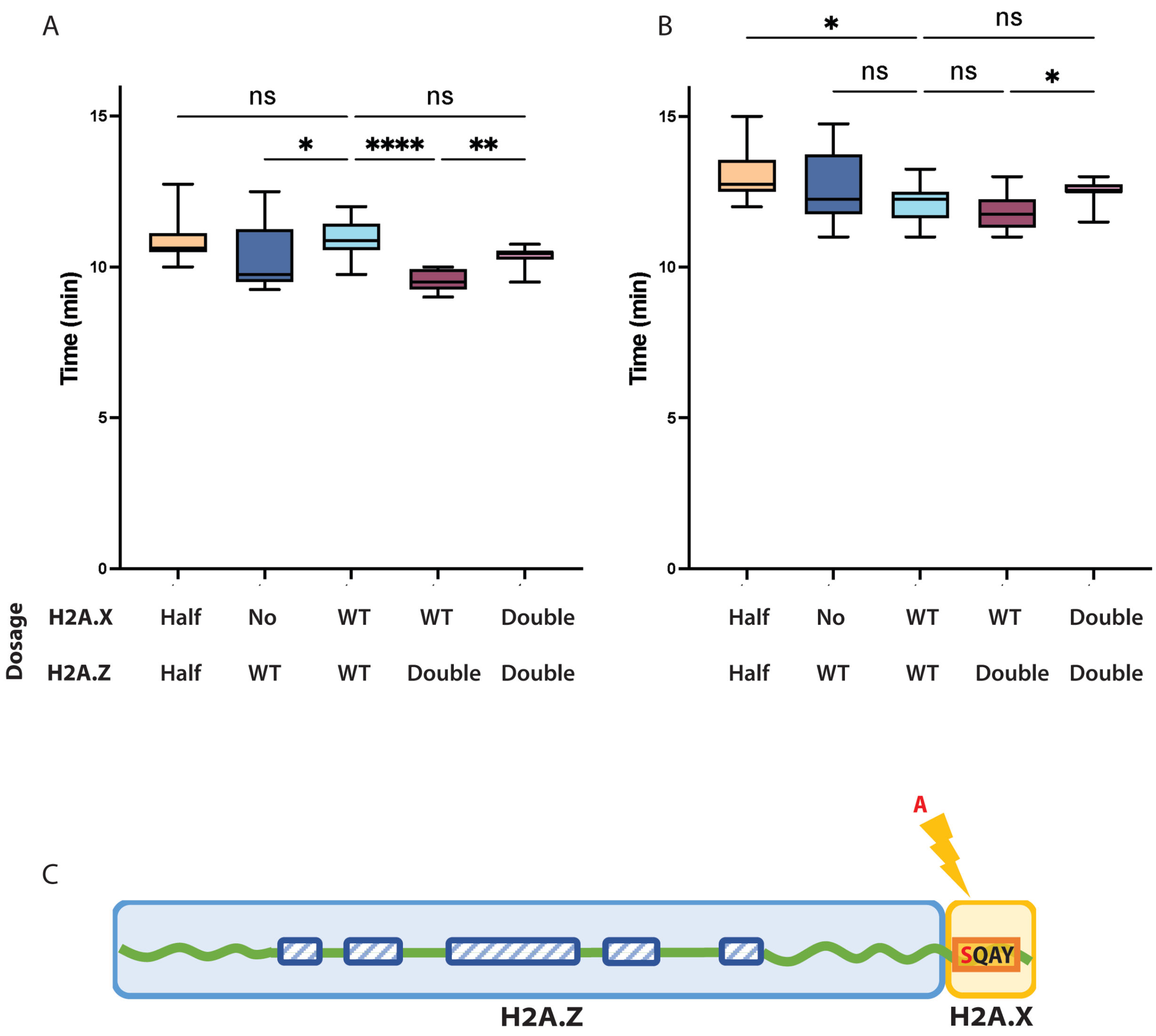

**Figure S4: Expression of many maternal and maternal-zygotic transcriptional units are altered in *Double H2Av* embryos**

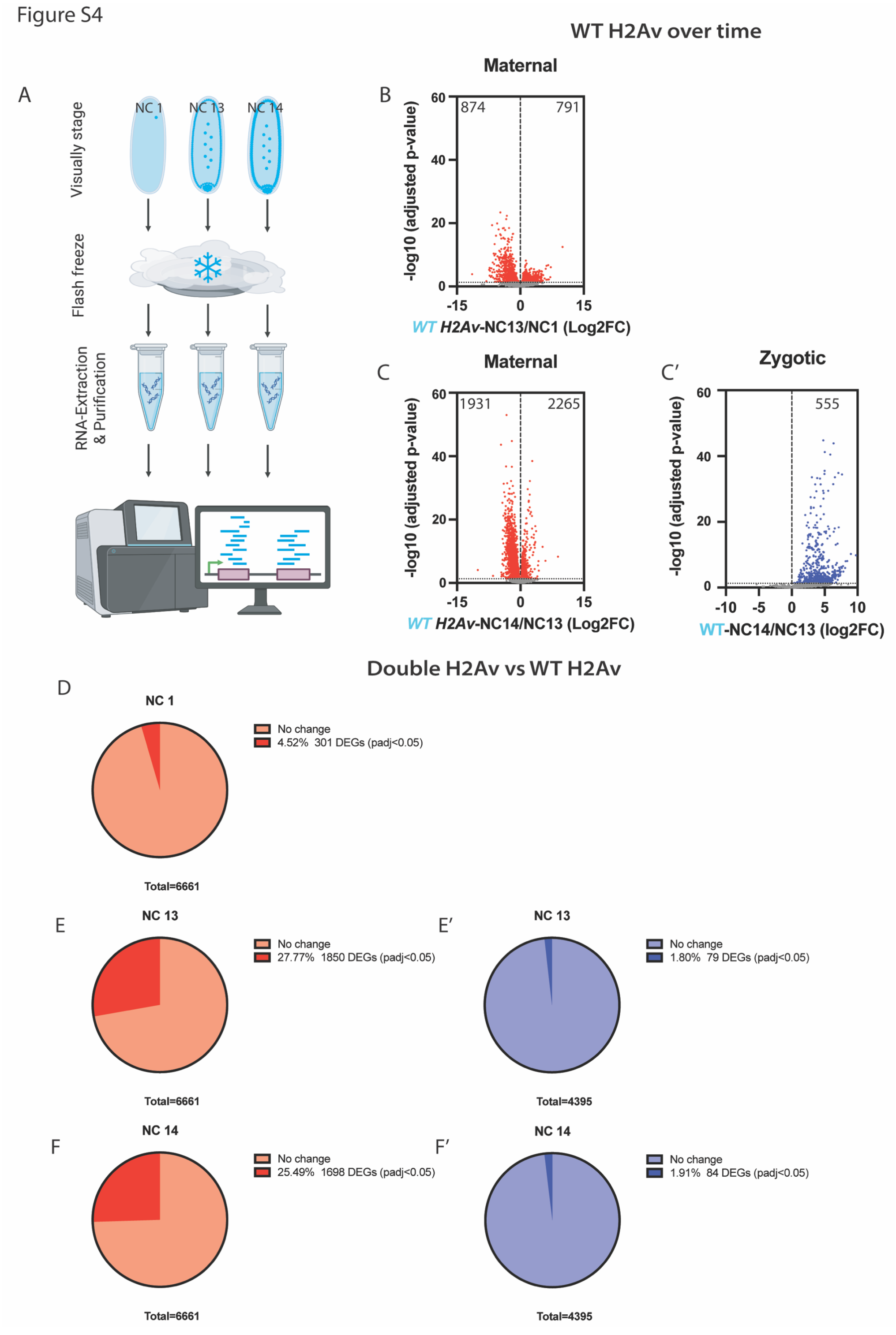

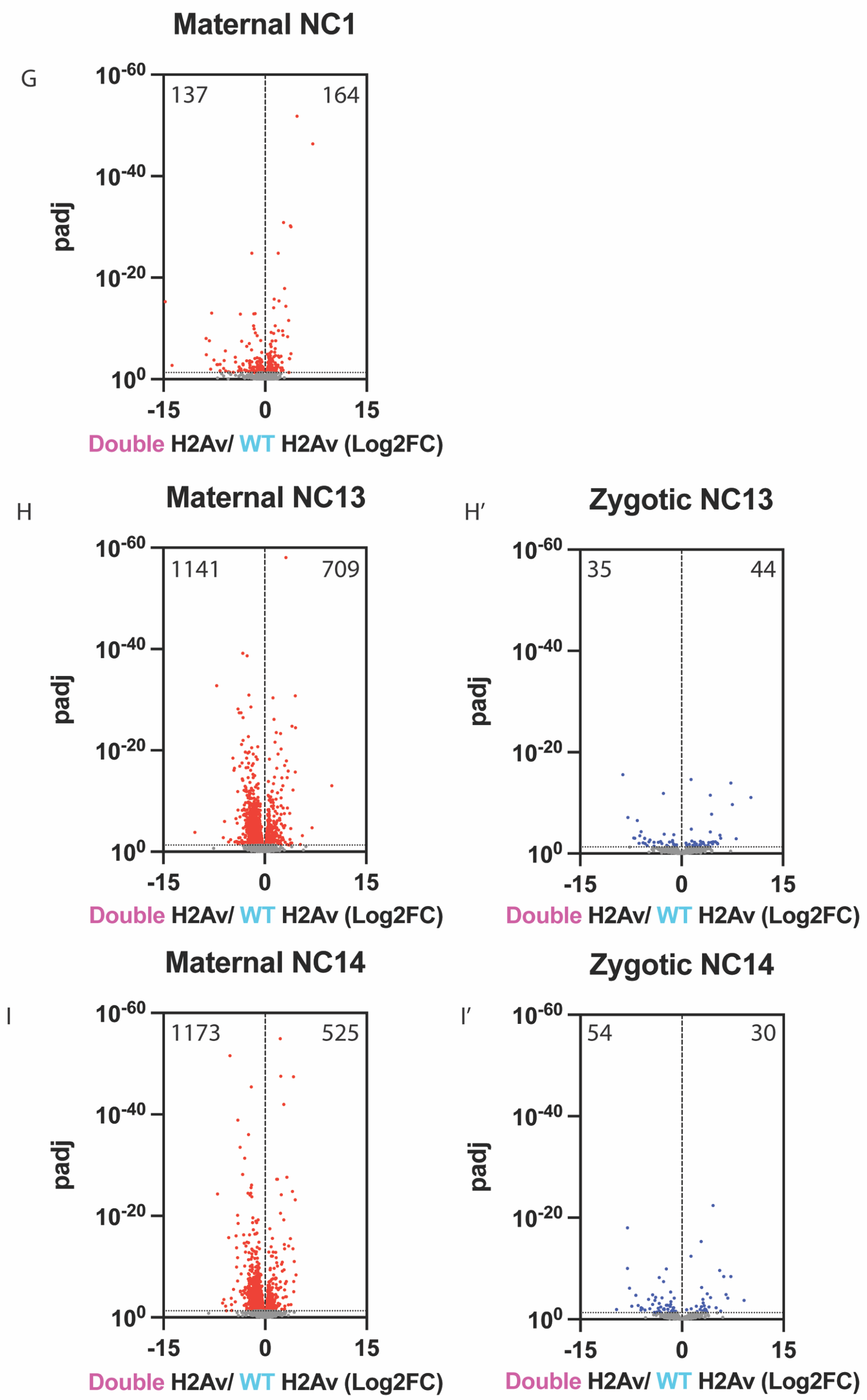
Maternal NC1
G
137
164
10-60
10-40
10-20
100
padj
-15
0
15
Double H2Av/ WT H2Av (Log2FC)
Maternal NC13
H
1141
709
Zygotic NC13
H′
35
44
Maternal NC14
I
1173
525
Zygotic NC14
I′
54
30

**Figure S5: Quadrants I and III are enriched in housekeeping genes**

Figure S5

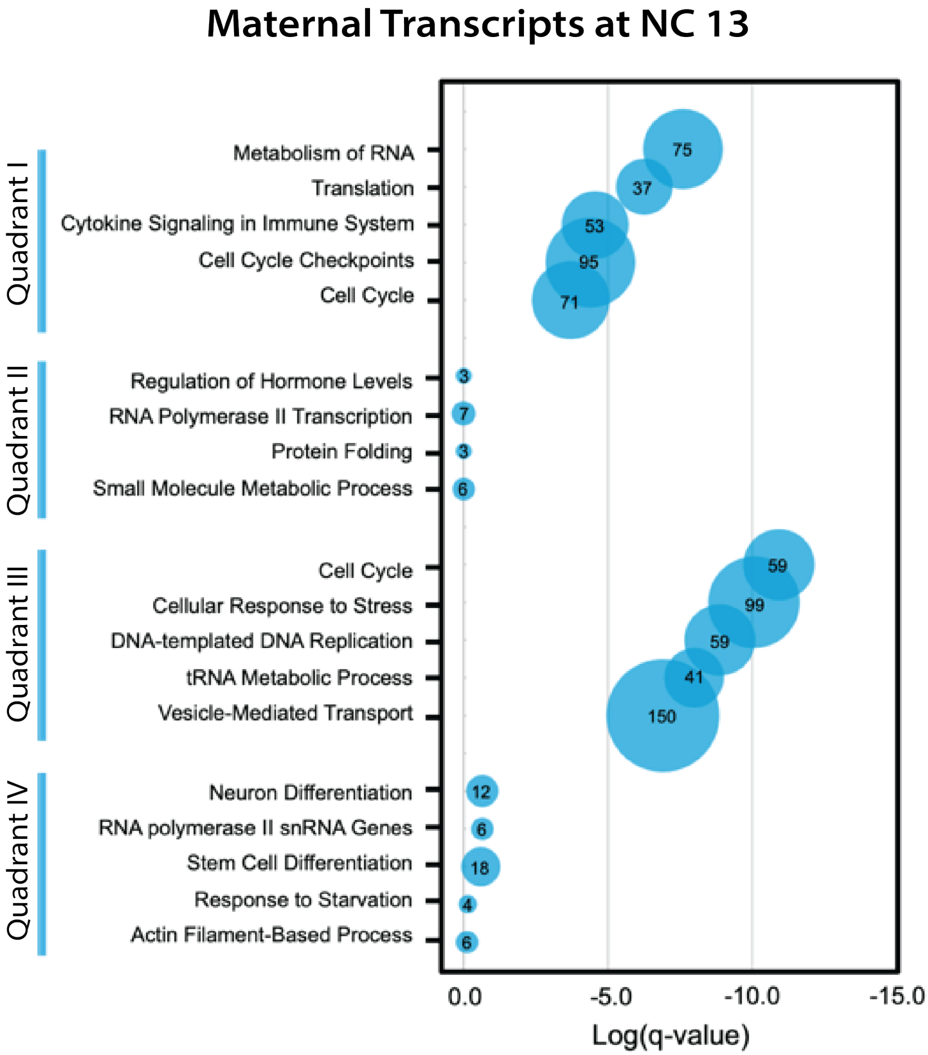

**Figure S6: *Domino* knockdown causes modest steady-state RNA changes but strong transcriptional disruption at ZGA**

Figure S6

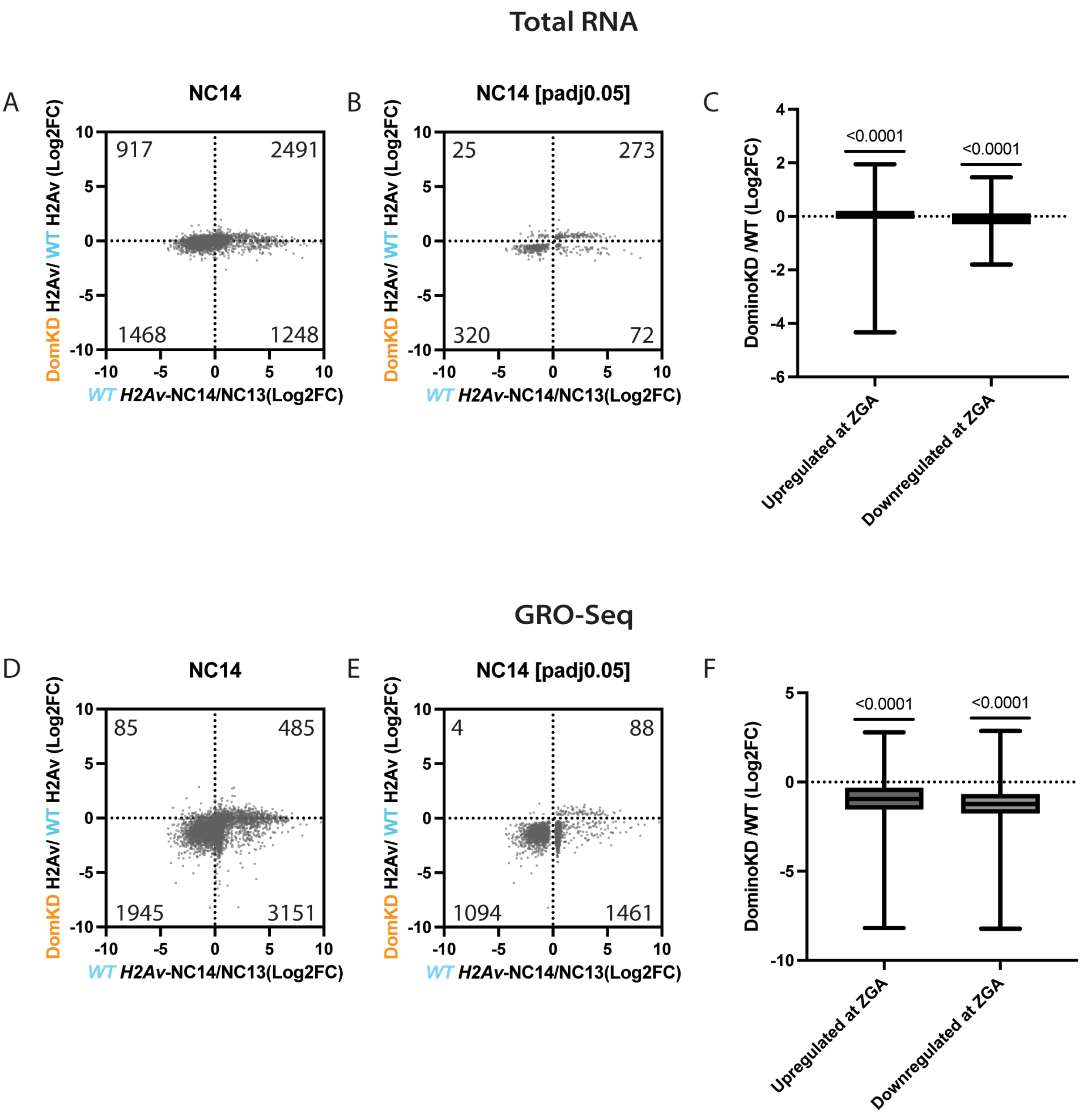

**Figure S7: NC 11 and NC12 timing in *Half Jabba* embryos**

Figure S7

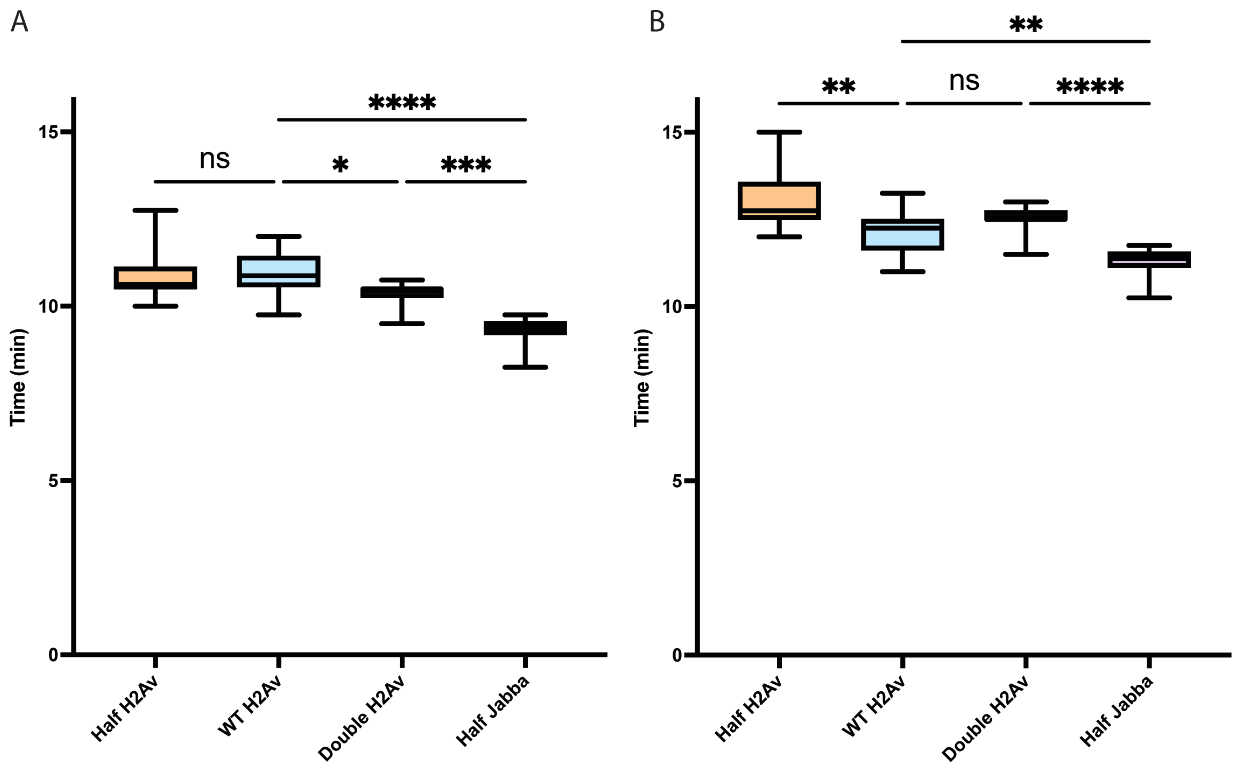

**Figure S8: *Half Jabba* embryos exhibit both expedited and delayed transcriptome remodeling**

Figure S8

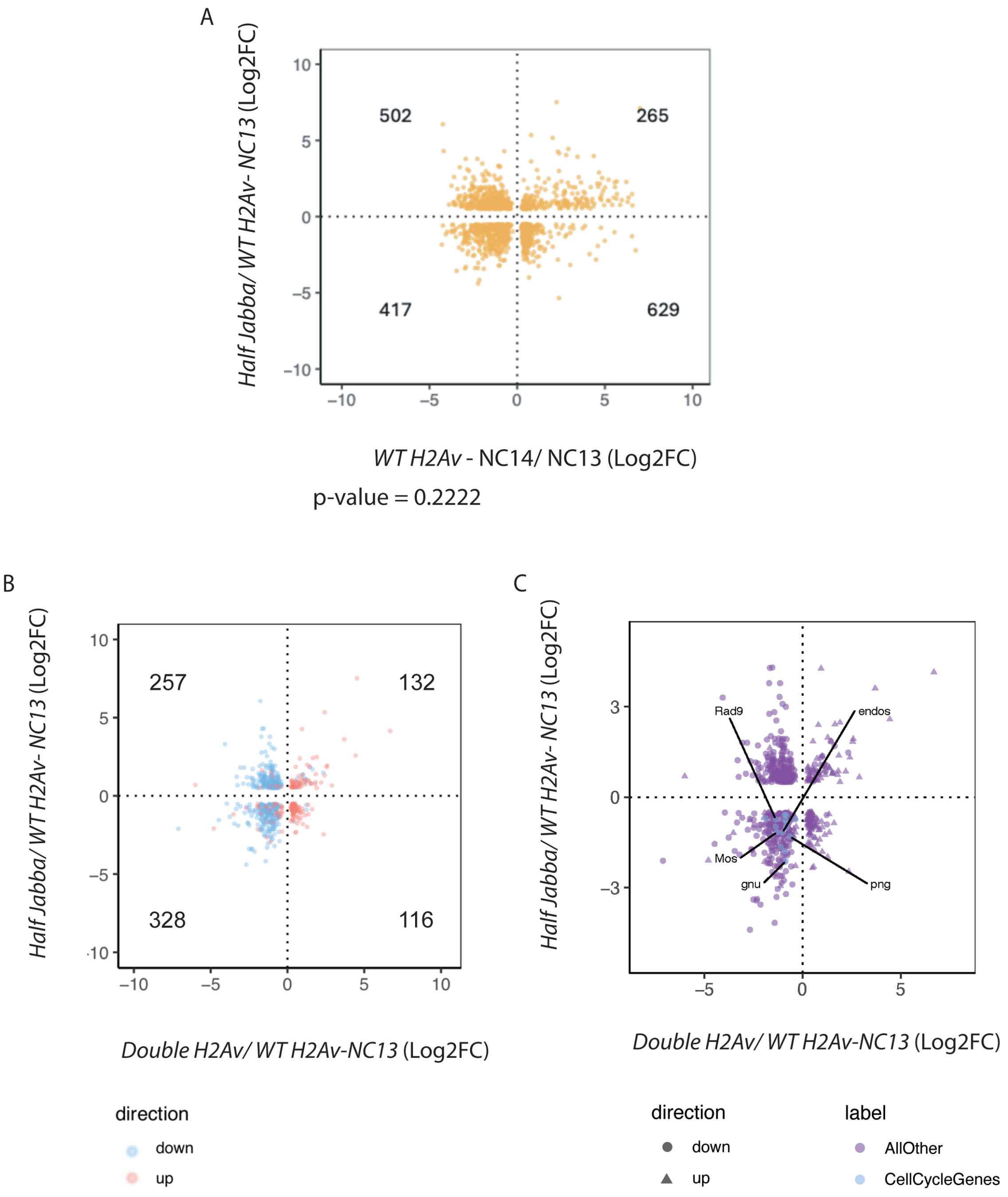

**Figure S9:** ***Double H2Av embryos*** **have increased levels of chromatin-associated H2Av**

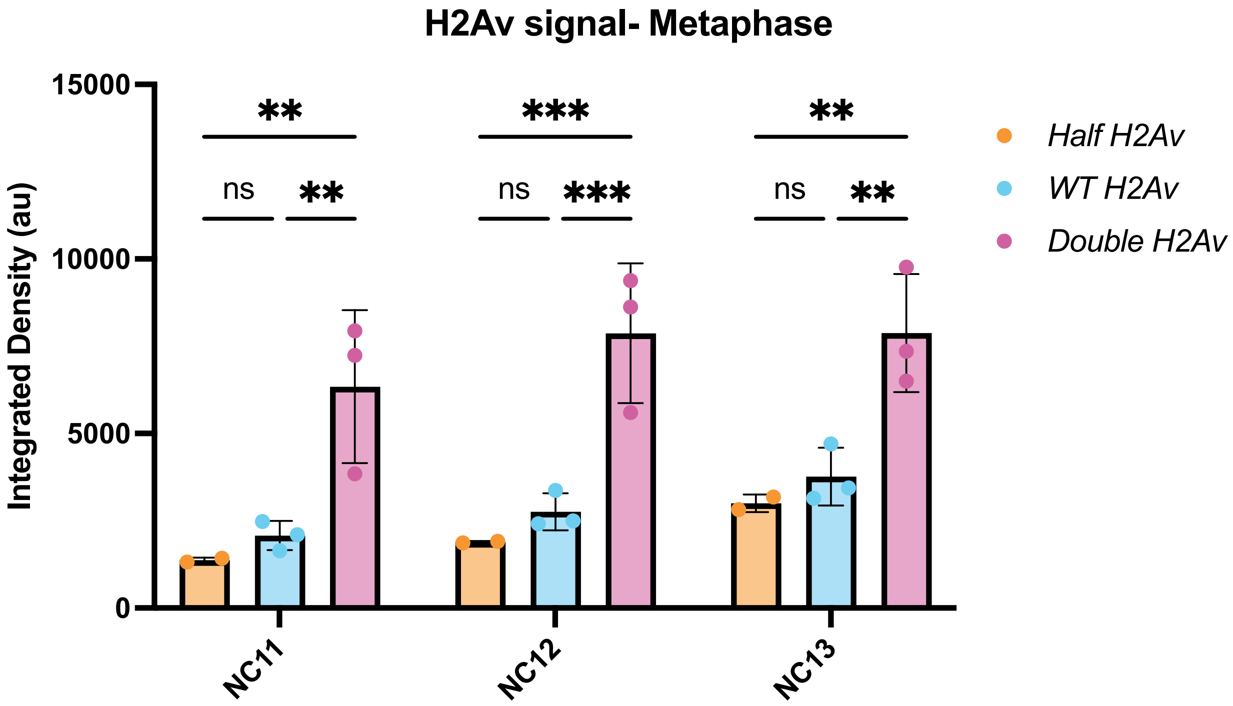

**Table S1 RNA-seq summary**

**Fly data**

| Sample Name | Condition | Genotype | Developmen | Paired or Single end | Batch |
|---|---|---|---|---|---|
| Egg_2X_1 | 2X_Egg | 2X-H2Av | Egg | SE | batch_1 |
| Egg_2X_2 | 2X_Egg | 2X-H2Av | Egg | SE | batch_1 |
| Egg_2X_3 | 2X_Egg | 2X-H2Av | Egg | SE | batch_1 |
| Egg_4X_1 | 4X_Egg | 4X-H2Av | Egg | SE | batch_1 |
| Egg_4X_2 | 4X_Egg | 4X-H2Av | Egg | SE | batch_1 |
| Egg_4X_3 | 4X_Egg | 4X-H2Av | Egg | SE | batch_1 |
| 2X-H2Av-NC13-1 | 2X_NC13 | 2X-H2Av | NC13 | PE | batch_2 |
| 2X-H2Av-NC13-2 | 2X_NC13 | 2X-H2Av | NC13 | PE | batch_2 |
| 2X-H2Av-NC13-3 | 2X_NC13 | 2X-H2Av | NC13 | PE | batch_2 |
| 2X-H2Av-NC14-1 | 2X_NC14 | 2X-H2Av | NC14 | PE | batch_2 |
| 2X-H2Av-NC14-2 | 2X_NC14 | 2X-H2Av | NC14 | PE | batch_2 |
| 2X-H2Av-NC14-3 | 2X_NC14 | 2X-H2Av | NC14 | PE | batch_2 |
| 4X-H2Av-NC13-1 | 4X_NC13 | 4X-H2Av | NC13 | PE | batch_2 |
| 4X-H2Av-NC13-2 | 4X_NC13 | 4X-H2Av | NC13 | PE | batch_2 |
| 4X-H2Av-NC14-1 | 4X_NC14 | 4X-H2Av | NC14 | PE | batch_2 |
| 4X-H2Av-NC14-2 | 4X_NC14 | 4X-H2Av | NC14 | PE | batch_2 |
| 4X-H2Av-NC14-3 | 4X_NC14 | 4X-H2Av | NC14 | PE | batch_2 |
| 1X-1-Egg | 1X_Egg | 1X-H2Av | Egg | PE | batch_3 |
| 1X-2-Egg | 1X_Egg | 1X-H2Av | Egg | PE | batch_3 |
| 1X-3-Egg | 1X_Egg | 1X-H2Av | Egg | PE | batch_3 |
| 1X-1-NC13 | 1X_NC13 | 1X-H2Av | NC13 | PE | batch_3 |
| 1X-2-NC13 | 1X_NC13 | 1X-H2Av | NC13 | PE | batch_3 |
| 1X-2-NC14 | 1X_NC14 | 1X-H2Av | NC14 | PE | batch_3 |
| 1X-3-NC14 | 1X_NC14 | 1X-H2Av | NC14 | PE | batch_3 |
| 2X-1-Egg | 2X_Egg | 2X-H2Av | Egg | PE | batch_3 |
| 1XDL_Egg_3 | 1XDL_Egg | 1X-Jabba | Egg | PE | batch_4 |
| 1XDL_Egg_combine | 1XDL_Egg | 1X-Jabba | Egg | PE | batch_4 |
| 1XDL_NC11_1 | 1XDL_NC11 | 1X-Jabba | NC11 | PE | batch_4 |
| 1XDL_NC11_combin | 1XDL_NC11 | 1X-Jabba | NC11 | PE | batch_4 |
| 1XDL_NC13_3 | 1XDL_NC13 | 1X-Jabba | NC13 | PE | batch_4 |
| 1XDL_NC13_combin | 1XDL_NC13 | 1X-Jabba | NC13 | PE | batch_4 |
| 1XDL_NC14_3 | 1XDL_NC14 | 1X-Jabba | NC14 | PE | batch_4 |
| 1XDL_NC14_combin | 1XDL_NC14 | 1X-Jabba | NC14 | PE | batch_4 |
| 1X_Egg_2_121024 | 1X_Egg | 1X-H2Av | Egg | PE | batch_4 |
| 1X_Egg_combine | 1X_Egg | 1X-H2Av | Egg | PE | batch_4 |
| 1X_NC11_3 | 1X_NC11 | 1X-H2Av | NC11 | PE | batch_4 |
| 1X_NC11_combine | 1X_NC11 | 1X-H2Av | NC11 | PE | batch_4 |
| 1X_NC13_3 | 1X_NC13 | 1X-H2Av | NC13 | PE | batch_4 |
| 1X_NC13_combine | 1X_NC13 | 1X-H2Av | NC13 | PE | batch_4 |
| 1X_NC14_1 | 1X_NC14 | 1X-H2Av | NC14 | PE | batch_4 |
| 1X_NC14_combine | 1X_NC14 | 1X-H2Av | NC14 | PE | batch_4 |

| | | | | | |
|---|---|---|---|---|---|
| 2X_Egg_1 | 2X_Egg | 2X-H2Av | Egg | PE | batch_4 |
| 2X_Egg_combine | 2X_Egg | 2X-H2Av | Egg | PE | batch_4 |
| 2X_NC11_2_121024 | 2X_NC11 | 2X-H2Av | NC11 | PE | batch_4 |
| 2X_NC11_combine | 2X_NC11 | 2X-H2Av | NC11 | PE | batch_4 |
| 2X_NC13_1_121024 | 2X_NC13 | 2X-H2Av | NC13 | PE | batch_4 |
| 2X_NC13_combine | 2X_NC13 | 2X-H2Av | NC13 | PE | batch_4 |
| 4X_Egg_1_121024 | 4X_Egg | 4X-H2Av | Egg | PE | batch_4 |
| 4X_Egg_combine | 4X_Egg | 4X-H2Av | Egg | PE | batch_4 |
| 4X_NC11_1_121024 | 4X_NC11 | 4X-H2Av | NC11 | PE | batch_4 |
| 4X_NC11_combine | 4X_NC11 | 4X-H2Av | NC11 | PE | batch_4 |
| 4X_NC13_2_121024 | 4X_NC13 | 4X-H2Av | NC13 | PE | batch_4 |

**Zebrafish data**

| Sample Name | Condition | Genotype | Developmen | Paired or Single end | Batch |
|---|---|---|---|---|---|
| GFP_2_5_4 | GFP_2_5 | H2A.Z OE | Early MBT | PE | batch_5 |
| GFP_2_5_5 | GFP_2_5 | H2A.Z OE | Early MBT | PE | batch_5 |
| GFP_2_5_6 | GFP_2_5 | H2A.Z OE | Early MBT | PE | batch_5 |
| GFP_4hpf_4 | GFP_4hpf | H2A.Z OE | Late MBT | PE | batch_5 |
| GFP_4hpf_5 | GFP_4hpf | H2A.Z OE | Late MBT | PE | batch_5 |
| GFP_4hpf_6 | GFP_4hpf | H2A.Z OE | Late MBT | PE | batch_5 |
| WT_2_5_1 | WT_2_5 | WT | Early MBT | PE | batch_5 |
| WT_2_5_2 | WT_2_5 | WT | Early MBT | PE | batch_5 |
| WT_2_5_3 | WT_2_5 | WT | Early MBT | PE | batch_5 |
| Wt_4hpf_1 | Wt_4hpf | WT | Late MBT | PE | batch_5 |
| Wt_4hpf_2 | Wt_4hpf | WT | Late MBT | PE | batch_5 |
| Wt_4hpf_3 | Wt_4hpf | WT | Late MBT | PE | batch_5 |
